# Radial glia promote microglial development through integrin α_V_β_8_-TGFβ1 signaling

**DOI:** 10.1101/2023.07.13.548459

**Authors:** Gabriel L. McKinsey, Nicolas Santander, Xiaoming Zhang, Kilian Kleemann, Lauren Tran, Aditya Katewa, Kaylynn Conant, Matthew Barraza, Kian Waddell, Carlos Lizama, Marie La Russa, Hyun Ji Koo, Hyunji Lee, Dibyanti Mukherjee, Helena Paidassi, E. S. Anton, Kamran Atabai, Dean Sheppard, Oleg Butovsky, Thomas D. Arnold

## Abstract

Microglia diversity emerges from interactions between intrinsic genetic programs and environment-derived signals, but how these processes unfold and interact in the developing brain remains unclear. Here, we show that radial glia-expressed integrin beta 8 (ITGB8) expressed in radial glia progenitors activates microglia-expressed TGFβ1, permitting microglial development. Domain-restricted deletion of *Itgb8* in these progenitors establishes complementary regions with developmentally arrested “dysmature” microglia that persist into adulthood. In the absence of autocrine TGFβ1 signaling, we find that microglia adopt a similar dysmature phenotype, leading to neuromotor symptoms almost identical to *Itgb8* mutant mice. In contrast, microglia lacking the TGFβ signal transducers *Smad2* and *Smad3* have a less polarized dysmature phenotype and correspondingly less severe neuromotor dysfunction. Finally, we show that non-canonical (Smad-independent) signaling partially suppresses disease and development associated gene expression, providing compelling evidence for the adoption of microglial developmental signaling pathways in the context of injury or disease.

## Introduction

A substantial body of evidence has shown that transforming growth factor beta (TGFβ) signaling is crucial for microglial development and function^1^. Most notably, *Tgfb1* null mice display severe neuropathological changes, including microgliosis, neuronal cell death, and synaptic loss^2,3^. However, these mice also have developmental brain vascular dysplasia and hemorrhage, making it difficult to distinguish primary versus secondary microglial changes. In line with a primary role for *Tgfb1*, culture of microglia with TGFβ induces microglia-specific gene expression and promotes microglial survival^2,4^. Furthermore, mice mutant for the microglial enriched gene *Lrrc33* (*Nrros*), which binds and presents TGFβ1 to its cognate receptor TGFβR2, have severe microglial defects, but no major vascular abnormalities^5,6^. Similarly, conditional deletion of *Tgfbr2* in microglia results in the disruption of microglial homeostasis, without affecting vascular or blood brain barrier function ^7–9^.

TGFβ1 is synthesized in a latent inactive complex that requires two key steps to be activated^10,11^. First, latent TGFβ1 is anchored to the surface of one cell type by a so-called “milieu” molecule, LRRC33 (NRROS). Second, the integrin α_V_β_8_ dimer on neighboring cells binds to and activates latent-TGFβ1, which can then signal to TGFβR2. Cryo-EM analyses of α_V_β_8_ in complex with TGFβ1 and LRRC33 support a model whereby the release and diffusion of active TGFβ ligand is not necessary for TGFβ signaling; rather, TGFβ1 is positioned by LRRC33 and α_V_β_8_ to interact with TGFβR2, reinforcing TGFβ signaling in the cell which expresses TGFβ1^10,11^. This “paracrine activation / autocrine signaling” model predicts that deletion of *Itgb8* in one cell type primarily effects a neighboring cell type which both presents and responds to TGFβ1. Consistent with this model, *Itgb8*, *Tgfb1*, *Tgfb1RGE*, *Lrrc33* mouse mutants have largely overlapping phenotypes^3,5,7,12^ However, the specific cell types expressing these various signaling components, and the timing of their interactions during brain development remain unclear.

In a previous study, we demonstrated a critical role for integrin α_V_β_8_ in presenting active TGFβ to microglia^7^. We found that in the absence of α_V_β_8_-mediated TGFβ signaling, microglia are developmentally arrested and hyper-reactive. Furthermore, the presence of these “dysmature” microglia (and not just the absence of mature microglia) causes astrocyte activation, loss of GABAergic interneurons, and abnormal myelination, aspects of pathology which underlie the development of a severe neuromotor syndrome characterized by seizures and spasticity. We found that this phenotype is entirely due to loss of TGFβ signaling in microglia during brain development because microglia-specific deletion of *Tgfbr2* during early stages of brain development completely recapitulated the *Itgb8* mutant phenotype.

In this study, we aimed to identify 1) The relevant *Itgb8* expressing cell types that mediate microglial TGFβ activation; 2) The developmental timing of *Itgb8-*mediated TGFβ signaling in microglia; 3) The cellular source and identity of the TGFβ ligand relevant for microglial development and homeostasis; 4) The relationship between developmentally disrupted microglia and disease associated microglia; and 5) The role of canonical (Smad-mediated) versus non-canonical TGFβ signaling in microglia. By systematically deleting *Itgb8* from different cell lineages of the adult and embryonic brain, we show that *Itgb8* expression, specifically in embryonic radial progenitors, is necessary for microglial differentiation, but is not required to maintain microglial homeostasis in adulthood. We find that microglia-expressed *Tgfb1* is required cell-autonomously for differentiation and, in contrast to *Itgb8*, is also required to maintain homeostasis. By analyzing various *Itgb8-TGFβ* pathway mutants, we find that non-canonical TGFβ signaling fine-tunes microglial maturation and homeostasis, such that upstream mutants have amplified microglia and neuromotor phenotypes compared to *Smad2/3* conditional mutants. These data support the model that ITGB8/TGFβ1 signaling via radial glia-microglial interactions is crucial for the embryonic maturation of microglia and that in the absence of this signaling, developmentally arrested dysmature microglia drive local neuroinflammation and disruptions in neurological function. Finally, we show that dysmature and disease associated microglia share transcriptional and epigenetic properties, pointing to core TGFβ-regulated gene networks active in both development and disease.

## Results

### Radial glia *Itgb8* expression promotes microglial development

We previously found that central nervous system (CNS)-wide deletion of *Itgb8* using *Nestin^Cre^* leads to microglial dysmaturation and associated neuromotor impairments nearly identical to those seen in mice with conditional deletion of *Tgfbr2* in microglia^7^. *Nestin^Cre^* constitutively removes *Itgb8* from all cells derived from the early neuroepithelium (astrocytes, oligodendrocytes, and neurons), and *Itgb8* is expressed in all of these cell types (Figures S1A and S1B)^13^. To determine which of these *Itgb8*-expressing cells are responsible for the phenotypes observed in *Itgb8^fl/fl^;Nestin^Cre^* mice, we generated six different *Itgb8* conditional knockout lines (*Itgb8^fl/fl^;Nestin^Cre^*, *Itgb8^fl/fl^;hGfap^Cre^*, *Itgb8^fl/fl^;Olig2^Cre^*, *Itgb8^fl/fl^;Syn1^Cre^*, *Itgb8^fl/fl^;Ald1l1^CreER-T^*^2^, *Itgb8^fl/fl^;Ubc^CreER-T^*^2^) to systematically delete *Itgb8* from astrocytes, oligodendrocytes or neurons individually and in combination (Figure S1C). As summarized in Figure S1, none of these cell-type specific *Itgb8* mutants produced the phenotypes observed in *Itgb8^fl/fl^;Nestin^Cre^* mice (motor dysfunction, microglial TMEM119/P2RY12 loss, GFAP upregulation). Most notably, conditional deletion of *Itgb8* using *hGFAP^Cre^* resulted in no obvious neuropathological changes despite recombining nearly all neurons, astrocytes and oligodendrocytes in the cortex (98.3+/−.3%, 98.1+/−2.7%, 100% respectively; Figures S1C-E). These data indicate that *Itgb8* is dispensable for maintaining microglial, neuro/glial and vascular homeostasis in adult mice.

To understand why *Nestin^Cre^* mediated *Itgb8* deletion resulted in a fulminant neuromotor phenotype, while deletion from all other brain cell types (alone or in combination) did not, we considered the unique features of this line ^13^. *Nestin^Cre^*-mediated recombination occurs at approximately embryonic day 10.5 (E10.5), resulting in loss of *Itgb8* in radial glia and their progeny; (Figure 1C-D, Figure S1). Interestingly, *hGFAP^Cre^* also recombines radial glia progenitors and their progeny, but three days later than *Nestin^Cre^* at ∼E13.5^14,15^. Both of these lines have recombination in almost all neurons and glia by adulthood (Figure S1). The sequential recombination of brain progenitors in *Itgb8^fl/fl^;Nestin^Cre^*(complete phenotype) and *Itgb8^fl/fl^;hGFAP^Cre^* (no apparent phenotype) suggests that the developmental timing of *Itgb8* expression in, or deletion from, early embryonic neuroepithelial progenitor cells or radial glia is responsible for the stark phenotypic differences in these mice.

**Figure 1.**
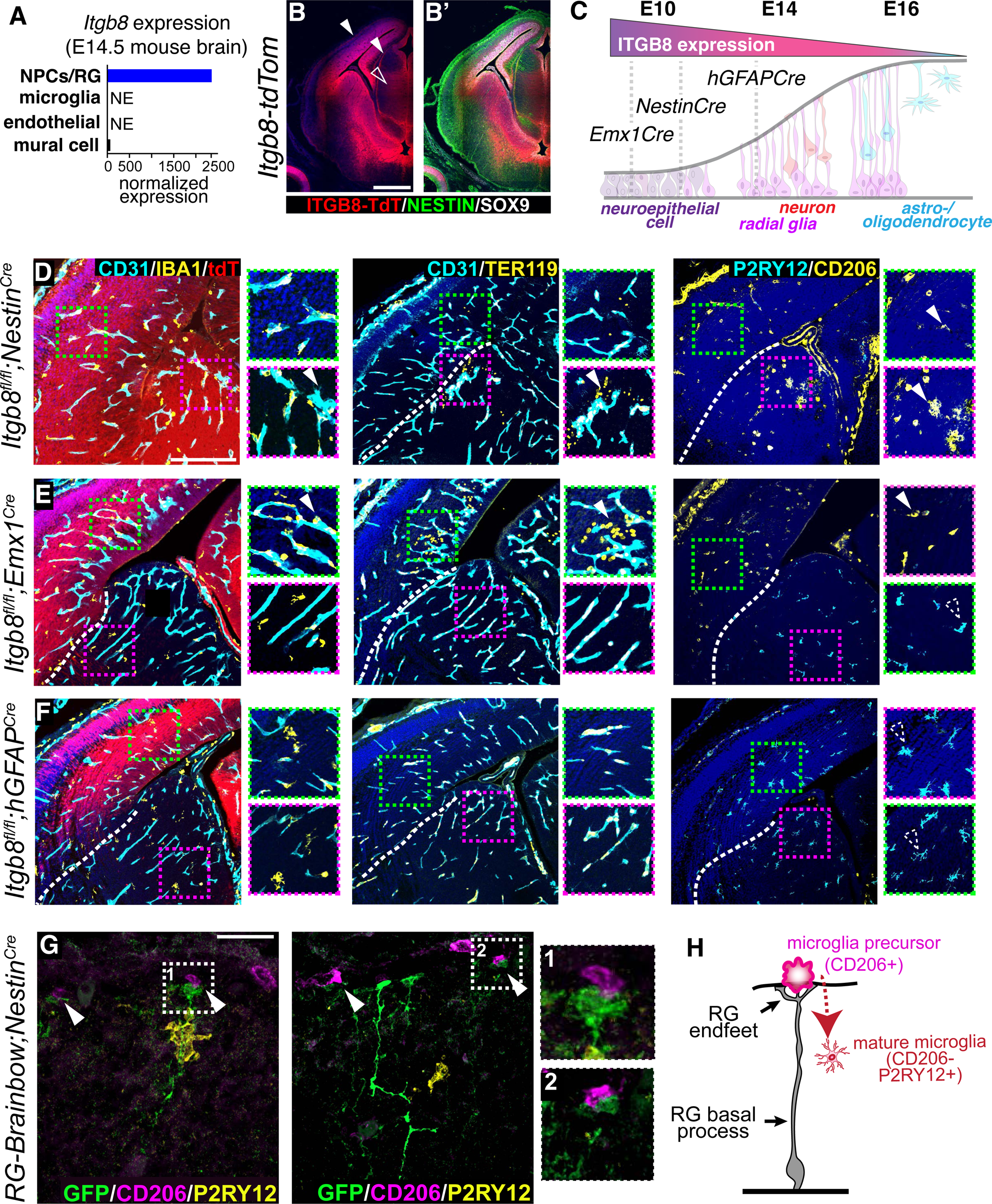
Deletion of *Itgb8* in early embryonic radial glia disrupts microglial maturation. **A)** Analysis of *Itgb8* expression in the E14.5 mouse embryo in neural progenitor cells (NPCs) and radial glia, microglia, endothelial cells, and mural cells^16^. **B)** *Itgb8^tdT^* reporter expression confirms strong *Itgb8* expression in SOX9+NESTIN+ radial progenitors at E14.5. Open arrowhead marks radial glia fibers; closed arrowhead marks ramified radial glia endfeet at the surface of the neuroepithelium. **C)** Model describing developmental expression of *Itgb8* in neuroepithelium and radial glia, and correlation with sequential timing of Cre recombination in *Emx1^Cre^*, *Nestin^Cre^*and *hGFAP^Cre^* lines. **D-F)** Deletion of *Itgb8* from neuroepithelial and radial progenitors using indicated Cre lines. Coronal brain sections stained for tdT (Cre recombination, red), vascular endothelium (CD31, cyan), and macrophages/microglia (IBA1, yellow); hemorrhage (red blood cells marked by TER119 (yellow) observed outside of vascular lumen (CD31, cyan); microglia precursors (CD206, yellow) and committed/homeostatic microglia (P2RY12, cyan). **G)** E14.5 brain section from *Emx1^Cre^;RG-brainbow* mouse stained for membranous GFP (individual recombined radial glia; endfeet, green), microglia precursors (CD206, magenta), and committed/homeostatic microglia (P2RY12, yellow). Arrowheads indicate foot process of radial glia contacting pial-associated CD206+ presumptive microglia precursor (model to right). Scale bar in B=500*μ*m, D=200*μ*m, G=25 *μ*m.

Published bulk and single cell RNA-Seq (Figures 1A and S2A)^16,17^, embryonic Flash-Tag ScRNA-seq (Figure S2B)^18^, and in situ hybridization (Figure S2C) show that *Itgb8* is expressed in neuroepithelial progenitor cells throughout the brain starting by E8.5, and that by E14.5 *Itgb8* is most highly expressed in radial glia progenitors, and less so in maturing astrocytes, oligodendrocytes and neurons^17,18^. We confirmed this using *Itgb8-IRES-tdTomato (Itgb8^tdT^)* reporter mice, finding intense tdT expression in ventricular zone progenitors at E14.5 (Figures 1B and S2D)^19^ as well as tdT expression in radial glia fibers and endfeet in the embryonic meninges (Figure S2E).

To more directly test the hypothesis that early developmental expression of *Itgb8* is necessary for microglial maturation, we analyzed *Itgb8* conditional knockouts (*Nestin^Cre^*, *hGFAP^Cre^*, *Olig2^Cre^, Syn1^Cre^*) at E14.5. We also generated *Itgb8^fl/fl^;Emx1^Cre^*mice, in which Cre is expressed in neocortical neuroepithelial cells just before the formation of radial glia at E9^20^ (Figure 1C). As we previously reported^7^, we found that *Itgb8^fl/fl^;Nestin^Cre^*mice develop vascular dysplasia and hemorrhage in the brain starting at E11.5 (not shown), which progresses in a ventral-dorsal fashion to include all periventricular vessels by E14.5 (Figure 1D). Coinciding with these vascular changes, we observed that IBA1+ macrophages were strongly associated with dysplastic vessels in *Itgb8^fl/fl^;Nestin^Cre^* mutants (IBA1/CD31 apposition in Figure 1D), and were characteristically dysmature, i.e. lacking expression of mature/homeostatic microglia marker P2RY12, and reciprocally upregulating (or persistently expressing) CD206, a marker of undifferentiated microglial precursors and border associated macrophages (BAMs) (arrowheads in right column of Figure 1D)^21,22,23,24^. While the vascular phenotype and hemorrhage was largely localized to areas near the cerebral ventricles (the periventricular vascular plexus, PVP), we observed microglia lacking P2RY12 staining throughout the brain, including areas far from hemorrhage, suggesting that the two phenotypes might be dissociable, i.e. that hemorrhage is not causing these widespread microglial changes or vice-versa.

Similar to *Itgb8^fl/fl^;Nestin^Cre^* embryos, *Itgb8^fl/fl^;Emx1^Cre^* E14.5 embryos had vascular abnormalities and hemorrhage near the cerebral ventricles, concomitant with microglial changes throughout the developing cortex and hippocampus. However, these vascular/hemorrhage and microglial abnormalities in *Itgb8^fl/fl^;Emx1^Cre^*mice were strikingly domain-specific; hemorrhage and microglial changes occurred only in the developing cerebral cortex and hippocampus, where Cre-recombinase had been active (as indicated by tdTomato (tdT) Cre-reporter expression), but not in the underlying striatum (Figure 1E). In sharp contrast, *Itgb8^fl/fl^;hGFAP^Cre^* mice lacked any apparent vascular or microglial phenotype (Figure 1F). Similar to *Itgb8^fl/fl^;hGFAP^Cre^* mice, we observed no obvious vascular or microglial phenotype in *Itgb8^fl/fl^;Olig2^Cre^* or *Itgb8^fl/fl^;Syn1^Cre^*mutants (Figure S3). These Cre lines are active at or before E14.5, but not in radial glia. Altogether, based on the timing of *Itgb8* expression in the brain and the timing of recombination in these various Cre lines (*Emx1^Cre^* ∼E9; *Nestin^Cre^* ∼E10.5; *hGFAP^Cre^* ∼E13.5, see Figure 1C), our data indicate that *Itgb8* expression in early stage radial glia progenitors promotes both vascular and microglia maturation, and that expression of *Itgb8* in astrocytes, oligodendrocytes, and in post-mitotic neurons is dispensable for brain vascular and microglia development, and for ongoing cellular homeostasis.

We sought to further examine the interactions between radial glia and microglia. Initial descriptions of microglia by del Rio-Hortega proposed that microglia are developmentally derived from the inner layer of the meninges, the pia mater^25^. While CD206 is a marker of borderzone macrophages in adulthood, recent genetic fate-mapping reports show that microglia are derived from a CD206 expressing undifferentiated macrophage precursor^24,22,23^. Upon differentiating, these progenitors down-regulate the expression of CD206 and up-regulate microglial markers such as P2RY12. To determine whether embryonic radial glia make contact with CD206+ microglial precursors, we crossed *Emx1^Cre^*mice to a mouse line (*RG-Brainbow*) that expresses a Cre-dependent plasma membrane-tagged fluorescent reporter in a mosaic fashion under the GLAST promotor, is expressed in radial glia^26^. We then examined the interaction between radial glial endfeet and CD206+ macrophages in the meninges, where most embryonic CNS-associated CD206+ macrophages reside. Analysis of the mosaic expression of mGFP (membranous) in E14.5 *Emx1^Cre^;RG-Brainbow* embryos revealed contact between radial glia endfeet and CD206+ macrophages in the innermost layer of the meninges (Figure 1G), consistent with the model that meningeal CD206+ microglial precursor interaction with *Itgb8* expressing radial glial endfeet is necessary for the TGFβ-dependent differentiation of these progenitors (Figure 1H). In total, these data suggest that *Itgb8* expression in early embryonic radial glia is necessary for the TGFβ-dependent differentiation of CD206+ microglial precursors, and that in the absence of *Itgb8*, these CD206+ precursors populate the brain, but fail to express microglial-specific markers.

### Domain-specific microglial dysmaturity persists in adult *Itgb8^fl/fl^;Emx1^Cre^* mice

We next followed *Itgb8^fl/fl^;Emx1^Cre^* mice into adulthood to see how developmental phenotypes might evolve over time. Similar to *Itgb8^fl/fl^;Nestin^Cre^*, mice, vascular defects and hemorrhage resolved, while microglial changes persisted. To examine the transcriptional properties of microglia in *Itgb8^fl/fl^;Emx1^Cre^*mice, we analyzed our transcriptomic data derived from fluorescence-activated cell sorting of *Itgb8^fl/fl^;Emx1^Cre^* cortical and hippocampal microglia (Yin et al. Nature Immunology, In Press). To explore our hypothesis that cortical and hippocampal microglia in *Itgb8^fl/fl^;Emx1^Cre^*microglia are developmentally blocked in an immature state, we compared these transcriptomic data to a recently published transcriptional analysis that identified 7 gene clusters associated with microglia at various stages of development, from an early yolk sac progenitor stage through embryogenesis and into adulthood^27^. When we compared these developmental gene clusters in adult *Itgb8^fl/fl^;Emx1^Cre^*dysmature microglia to adult control microglia, we found a striking enrichment for genes in clusters 3-5, which represent transcripts found in yolk sac-derived microglial precursors and early embryonic microglia (Figures 2A and 2B). Conversely, genes that were enriched in clusters 6 and 7, which represent late embryonic/neonatal and adult stages were strongly downregulated (Figure 2B). Immunohistochemical analysis confirmed that the dysmature microglial phenotype was characterized by loss of homeostatic microglial markers and upregulation of reactive/immature markers (P2RY12^neg^, TMEM119^neg^; CD206^hi^, CLEC7a^hi^ and APOE^hi^) and was tightly restricted to the *Emx1^Cre^* recombination domain (Figures 2C-G). Outside this domain, microglia were comparable to control mice. Surprisingly, and in contrast to *Itgb8^fl/fl^;Nestin^Cre^*mice, *Itgb8^fl/fl^;Emx1^Cre^ mice* had no obvious neuromotor deficits or early mortality, suggesting that the profound motor disturbances in *Itgb8^fl/fl^;Nestin^Cre^* mice are not solely derived from microglial changes in the motor or sensory cortex.

**Figure 2.**
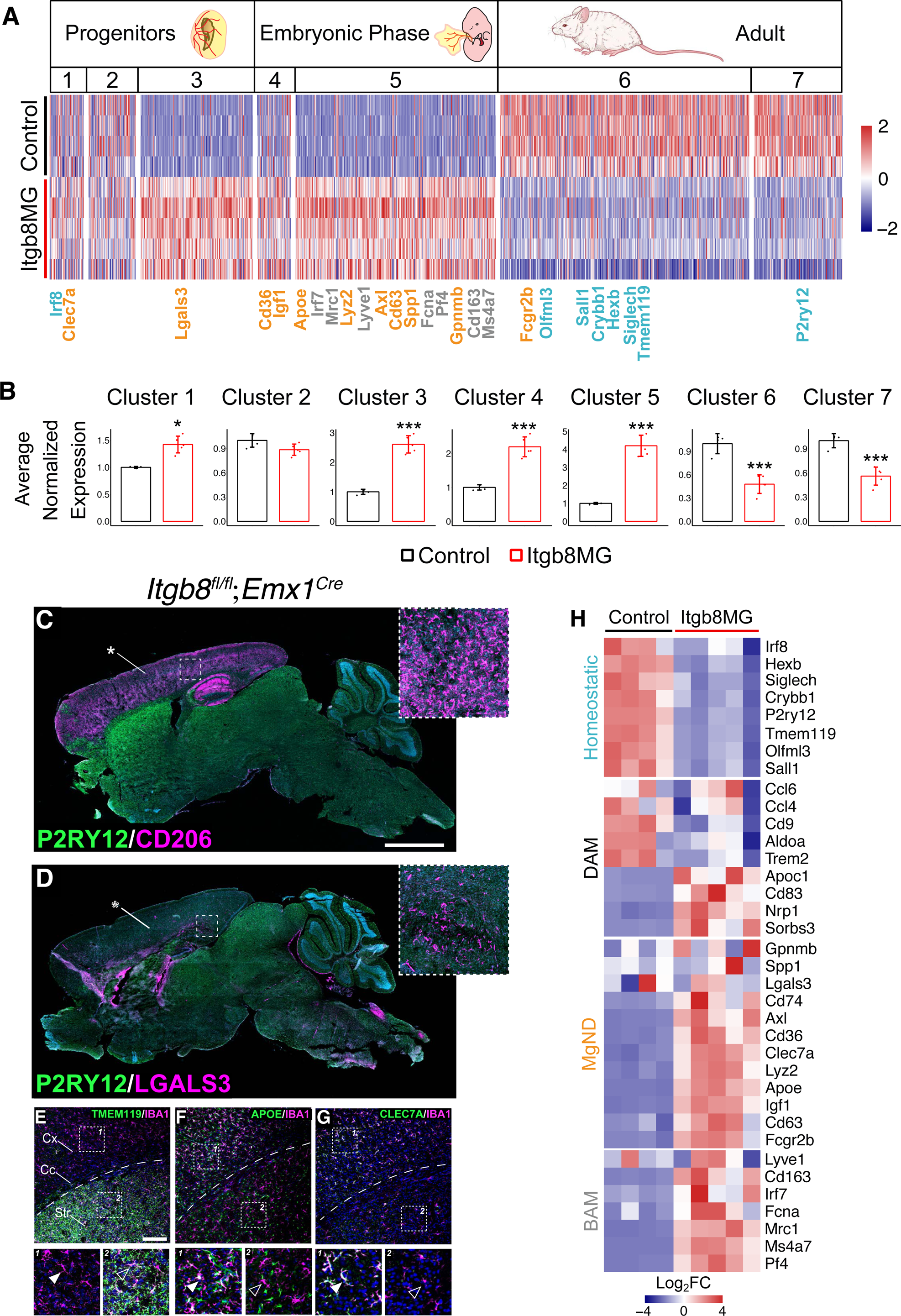
*Emx1-Cre* deletion of *Itgb8* results in anatomically restricted blockage of microglial differentiation. **A)** Comparison of the transcriptional properties of adult *Itgb8^fl/fl^; Emx1^Cre^* mutant and control microglia to stage specific developmental markers reveals that dysmature microglia retain the gene expression profiles of early embryonic microglia.^27^ **B)** Analysis of developmental gene cluster expression reveals enrichment for progenitor (cluster 1) and early embryonic phase (clusters 3-5) enriched gene sets. **C)** Whole brain sagittal immunostaining of adult *Itgb8^fl/fl^; Emx1^Cre^* mice revealed anatomically restricted maintenance of the microglial precursor marker CD206 in the cortex and hippocampus (asterisk), accompanied by loss of the homeostatic marker P2RY12. **D)** Increased expression of the MGnD marker LGALS3 in a subset of cortical and hippocampal microglia in *Itgb8^fl/fl^; Emx1^Cre^* mice (asterisk). Closed and open arrowheads in **E-G)** mark cortical and striatal microglia respectively. **E)** Downregulation of the microglial homeostatic marker TMEM119 in the cortex of a *Itgb8^fl/fl^; Emx1^Cre^* mouse. **F)** Cortex-restricted upregulation of the microglial reactive marker APOE in IBA1+ cells of the cortex (green cells). **G)** Cortex-restricted upregulation of the microglial reactive marker CLEC7a. Cx= cerebral cortex; Cc= corpus callosum; Str= striatum; Dashed line= cortical/striatal boundary. Scale bar in C= 2mm, E=150*μ*m.

Interestingly, we found that many of the genes enriched in dysmature *Itgb8^fl/fl^;Emx1^Cre^* microglia are similarly enriched in various disease and injury states (DAM, MGnD)^2,28^ (Figures 2A, 2H). Furthermore, many of these disease and injury-associated genes are developmentally expressed (Figure 2A), supporting the model that disease and injury-associated microglia recapitulate transcriptional states associated with progenitor and early embryonic developmental stages^29,30^. In addition to these disease and injury associated genes, dysmature *Itgb8^fl/fl^;Emx1^Cre^*microglia expressed BAM markers (Figures 2A and 2H), which are also expressed developmentally in microglia. Together, these experiments indicate that domain-specific deletion of *Itgb8* in radial glia progenitors leads to a zonally restricted arrest in microglial development. These developmentally arrested microglia appear to remain trapped in both time (an early embryonic progenitor state) and space (bounded by progenitor Cre recombination domains), and share transcriptional properties with disease and injury-associated microglia and borderzone macrophages.

### Dysmature microglia show pervasive epigenetic changes associated with MGnD and BAM gene upregulation

To better define the mechanistic basis for the transcriptional changes in *Itgb8* mutant dysmature microglia, we performed assays for transposase-accessible chromatin with sequencing (ATAC-seq) to identify regions of differentially accessible chromatin in *Itgb8^fl/fl^;Emx1^Cre^*dysmature microglia versus microglia from control brains. We also performed ChIP-Seq assays of histone H3 lysine 9 acetylation (H3K9ac) to identify the changes in active promoters and enhancers in *Itgb8^fl/fl^;Emx1^Cre^* dysmature microglia. We found a moderate linear correlation between genes associated with ATAC-seq defined differentially accessible peaks (DAPs) and *Itgb8^fl/fl^;Emx1^Cre^*dysmature microglia gene expression (R=.42, p<2.2e-16)(Figure 3B) and stronger correlation between H3K9ac DAPs in dysmature microglia and changes in dysmature microglia gene expression (R=.82, p<2.2e-16)(Figure 3C). Of the genes with the most significantly reduced levels of open chromatin and H3K9ac enrichment, many were markers of microglial identity (*P2ry12*, *HexB*, *Sall1*) (Figures 3A-3F and S4A). In contrast, genes with the most significantly increased levels of open chromatin and H3K9ac enrichment were largely BAM (*Mrc1*, *Pf4*, *Ms4a7*) or MGnD markers (*ApoE*, *Clec7a*, *Axl*) (Figures 3, S4B and S4C). Many of these microglia identity genes are directly regulated by SMAD4 in wild type microglia, while many BAM and MGnD genes are ectopically bound by SMAD4 in mice lacking *Sall1* expression in microglia, consistent with direct SMAD4-mediated gene activation or repression initiated by ITGB8/TGFβ signaling^31^. De novo motif enrichment analysis of DAPs most highly increased or decreased in ATAC-seq or H3K9ac-seq recovered motifs recognized by PU.1, MEF2C, RUNX1, SMAD2, and MAFB, all of which are transcription factors known to be relevant for microglial development and homeostasis^1,32–34^ (Figure 3F). The changes in the availability of these gene regulatory elements in *Itgb8^fl/fl^;Emx1^Cre^*cortical and hippocampal microglia are therefore likely to reflect widespread changes in the binding and regulatory activity of transcription factors that are crucial for the acquisition and maintenance of microglial identity.

**Figure 3.**
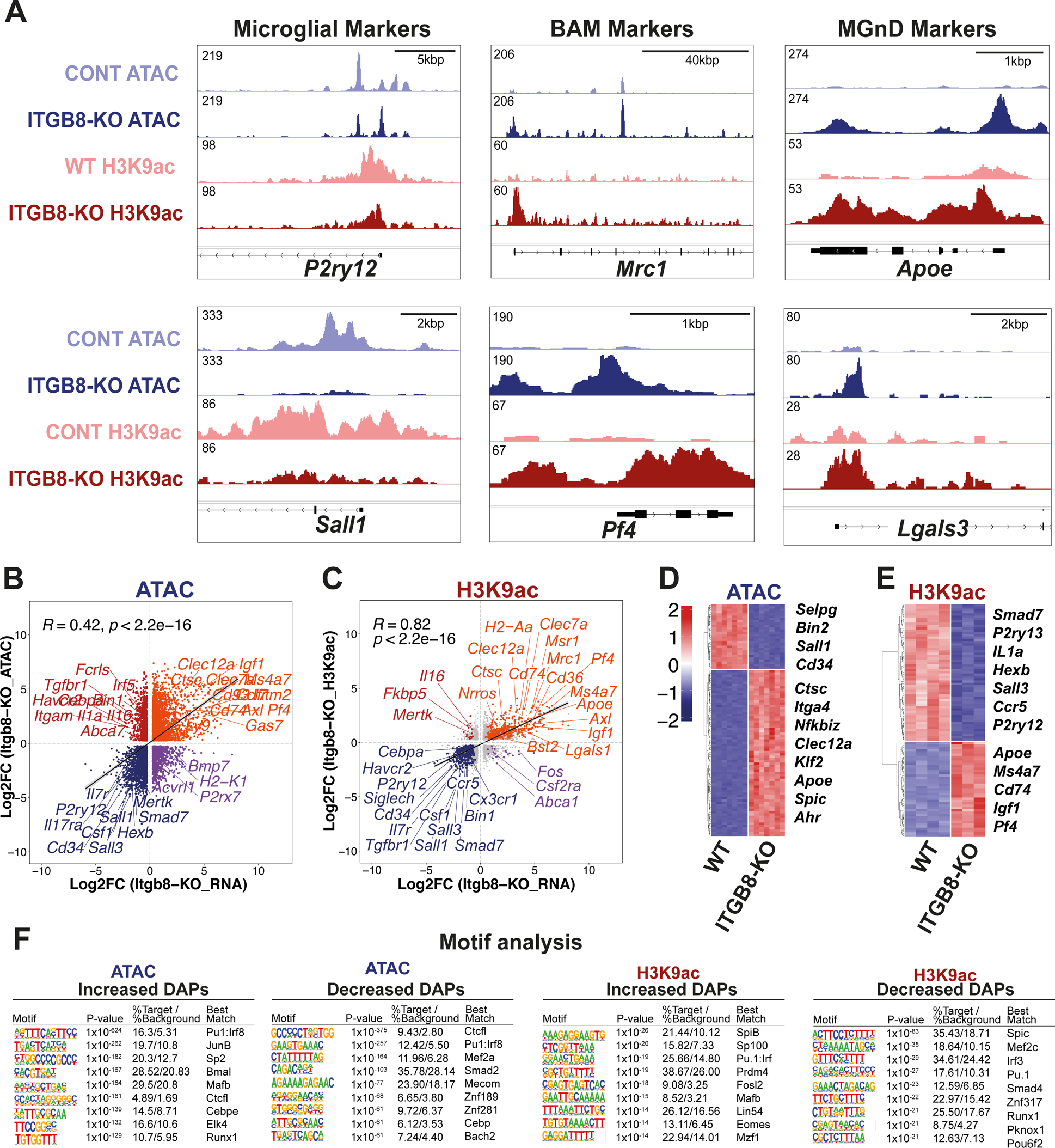
Dysmature microglia show pervasive epigenetic changes associated with MGnD and BAM gene upregulation. **A)** Tracks of ATAC-seq and H3K9ac ChIP-seq from *Itgb8^fl/fl^; Emx1^Cre^* microglia illustrating key homeostatic, border associated macrophage (BAM) and MgND marker gene bodies. **B)** Comparison between DEGs (RNA-seq) and DEPs (ATAC-seq) in cortical and hippocampal microglia from *Itgb8^fl/fl^; Emx1^Cre^* vs control brains (Padj < 0.05). Linear correlation between overlapping DEGs and DEPs, Log2FC. **C)** Comparison between DEGs (RNA-seq) and DEPs (H3K9ac ChIP-seq) in *Itgb8^fl/fl^; Emx1^Cre^* (Padj < 0.05) vs control brains. Linear correlation between overlapping DEPs and DEGs Log2FC. **D)** Heatmap of ATAC-seq showing top 100 Differentially accessible peaks (DEPs) comparing *Itgb8^fl/fl^; Emx1^Cre^* to control microglia (n=6, Padj < 0.05). **E)** Heatmap of H3K9ac ChIP-seq showing top 100 DEGs comparing *Itgb8^fl/fl^; Emx1^Cre^* to WT microglia. Padj < 0.05). **F)** Motif enrichment (HOMER) for positive or negative enriched peaks.

### *Tgfb1* is required cell-autonomously for microglial development

The only known biological function of ITGB8 is to activate TGFβ1 and TGFβ3^10,11^. Further supporting this, *Tgfb1^−/−^*mutant mice display neuropathology that closely resembles the various neurovascular and neuroimmune phenotypes observed in *Nestin^Cre^* and *Emx1^Cre^;Itgb8^fl/fl^* conditional mutants^3^. Recent structural biology studies^10,11^ reveal a mechanism of TGFβ1 activation where mature TGFβ1 signals within the confines of latent-TGFβ1; the release and diffusion of TGFβ1 is apparently not required. This “paracrine activation / autocrine signaling” model predicts that deletion of *Itgb8* on one cell type primarily affects another, and that deletion of *Tgfb1* will only affect the cells from which it is deleted. Here, we looked to test this model as it applies to microglial development.

To determine the developmental expression of *Tgfb1* in the brain, we examined a published RNAseq dataset, which showed strong expression of *Tgfb1* in brain macrophages, and weaker expression in brain blood vessels (both endothelial cells and PDGFRβ+ mural cells) at E14.5 (Figure 4A)^35^. Within the myeloid compartment, published bulk RNAseq from sorted microglia and BAMs^21^ indicates that *Tgfb1* is highly expressed in both microglia and BAMs throughout development (Figure 4B). In contrast, *P2ry12* and *Pf4* are specific markers for these cell types, respectively, and are expressed at early developmental stages. To test the hypothesis that cell-autonomous TGFB1 is necessary for microglial development, we generated *Tgfb1^fl/fl^;Cx3cr1^CreER^* (Figure 1D) and flox/null *Tgfb1^fl/GFP^;Cx3cr1^CreER^* (not shown) E14.5 embryos from dams treated with daily doses of tamoxifen for the three days prior to collection (E11.5-E13.5)^36,37^. *Tgfb1* RNAScope showed strong expression of *Tgfb1* in IB4 (Isolectin B4)+ blood vessels, IB4+ microglia and BAMs (meningeal and choroid plexus macrophages), and loss of *Tgfb1* expression in microglia and macrophages in *Tgfb1^fl/fl^;Cx3cr1^CreER^*embryos (Figures 4C and 4D). To specifically delete *Tgfb1* in BAMs or microglia we relied on our previous findings that *Pf4^Cre^* and *P2ry12^CreER^*specifically drive recombination in these populations respectively^24^. After finding that *P2ry12^CreER^* was unable to efficiently delete *Tgfb1* in all microglia, we sought to improve deletion efficiency using homozygous *P2ry12^CreER^*mice. We generated *Tgfb1^fl/GFP^;P2ry12^CreER/CreER^;Ai14* and *Tgfb1^fl/fl^;P2ry12^CreER/CreER^;Ai14* E14.5 embryos, inducing three days prior as with *Cx3cr1^CreER^* mice. Importantly, *P2ry12^CreER^* is a P2A fusion knock-in allele that retains P2RY12 protein expression in homo and heterozygous mice^24^. In contrast to *P2ry12-CreER* heterozygotes, which showed a partially penetrant phenotype, *Tgfb1^fl/GFP^;P2ry12^CreER/CreER^; Ai14* (Figure 4E) and *Tgfb1^fl/fl^;P2ry12^CreER/CreER^; Ai14* (not shown) embryos had a complete loss of P2RY12 expression that was identical to *Cx3cr1^CreER^*-mediated *Tgfb1* deletion, although meningeal macrophages were not recombined (Figure 4F). These data suggest that the microglial phenotype observed in these models is due to loss of *Tgfb1* in microglia, and not microglia progenitors or BAMs. To more directly test the function of *Tgfb1* in BAMs, we generated *Tgfb1^f/GFP^;Pf4^Cre^; Ai14* E14.5 embryos. We previously showed that *Pf4^Cre^* specifically recombines all BAMs including choroid plexus, meningeal and perivascular macrophages^24^. Of note, *Pf4* is highly expressed in yolk sac myeloid precursors^38^. Retention of recombined tdT+ cells in border-associated spaces, and lack of recombination in microglia, indicate that *Pf4* is an early marker of BAM-committed cells. Consistent with this, we observed specific recombination of meningeal and choroid plexus macrophages in E14.5 control and *Tgfb1* conditional knockouts, but no changes in microglial P2RY12 expression (Figures 4G and H). Together, these data indicate that microglia require self-produced TGFβ1 for their differentiation.

**Figure 4.**
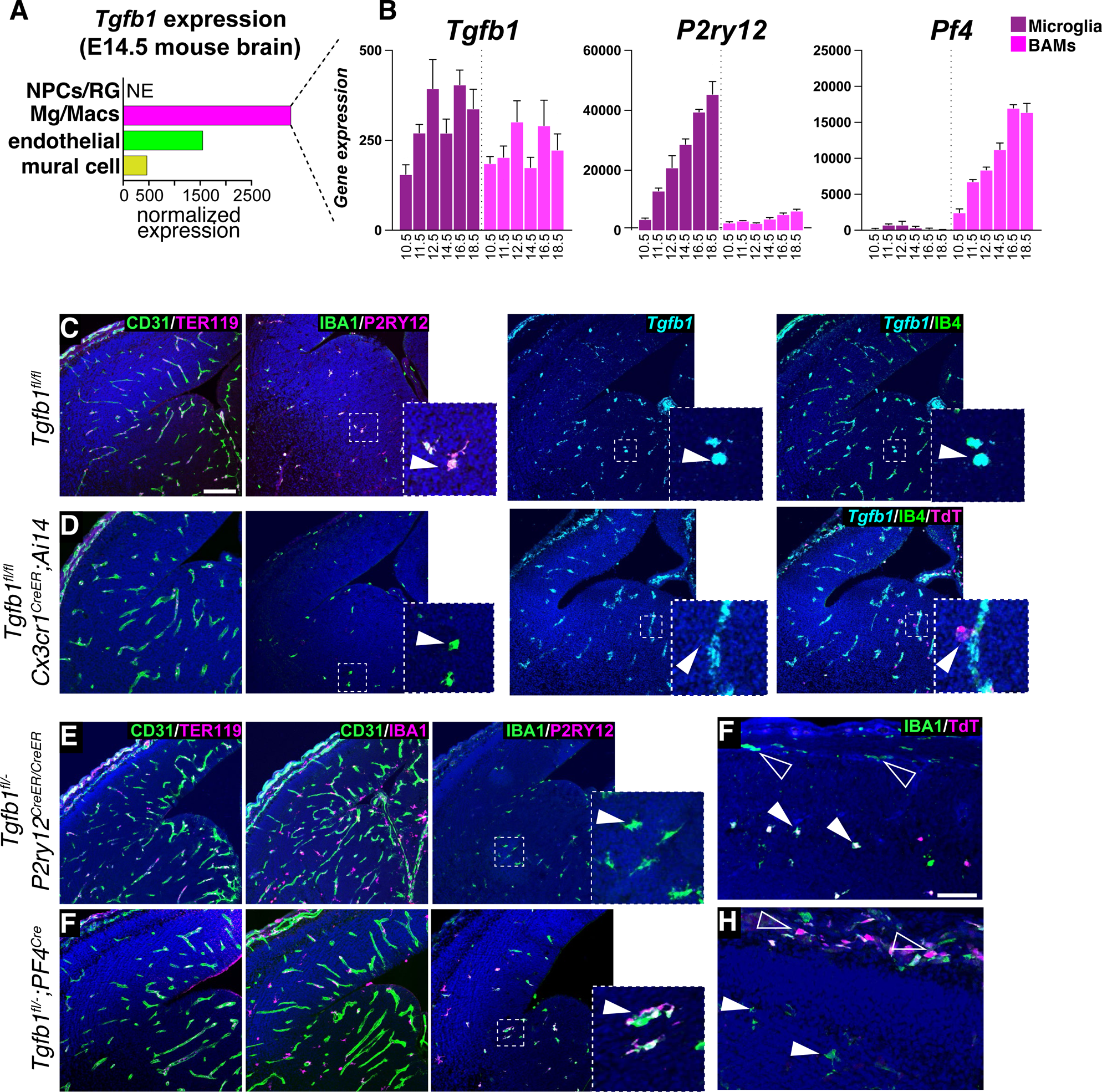
Microglia provide their own TGFβ1 to promote and maintain homeostasis. **A)** Sorted bulk-Seq analysis^21^ of embryonic microglia and BAMs reveals that *Tgfb1* is expressed in both microglia and BAMs during embryonic development, whereas **B)** *P2ry12* and *Pf4* are specific markers of these two respective populations. **C,D)** Analysis of control **(C)** and conditional *Cx3cr1^CreER^* mediated deletion (**D)** of *Tgfb1* deletion in the E14.5 forebrain following E11.5, 12.5 and 13.5 tamoxifen induction. Analysis revealed no hemorrhage (CD31 in green, TER119 in magenta), no change in macrophage/blood vessel association (IBA1 in magenta, CD31 in green), loss of the homeostatic marker P2RY12 (in magenta, IBA1 in green), and loss of *Tgfb1* (cyan) in Isolectin B4 (green) and Tdt (red) labeled microglia, but not in IB4 + labeled blood vessels. **E)** Analysis of conditional *P2ry12^CreER^* mediated deletion of *Tgfb1* deletion in E14.5 microglia the following E11.5, 12.5 and 13.5 tamoxifen induction. Analysis revealed no brain hemorrhage (CD31 in green, TER119 in magenta), no change in macrophage/blood vessel association (IBA1 in magenta, CD31 in green), and loss of the homeostatic marker P2RY12 (magenta), in IBA1+ (green) microglia. **F)** *P2ry12^CreER^* recombination, as shown by *ROSA-TDT (Ai14)* Cre reporter expression, was restricted to microglia (closed arrowheads), and was rarely seen in the overlying meninges (open arrowheads). **G)** Analysis of conditional *Pf4^Cre^* mediated deletion of *Tgfb1* deletion in the E14.5 forebrain. Analysis revealed no brain hemorrhage (CD31 in green, TER119 in magenta), no change in macrophage/blood vessel association (IBA1 in magenta, CD31 in green), and no loss of the homeostatic marker P2RY12 (magenta), in IBA1+ (green) microglia. H) *Pf4-Cre* recombination, as shown by *ROSA-TDT (Ai14)* Cre reporter expression, was restricted to the embryonic meninges (open arrowheads), and was not seen in microglia (open arrowheads). Scale bar in A=150*μ*m, F-50*μ*m.

We next looked to determine the source of *Tgfb1* for brain vascular development, and to probe potential interactions between developing brain blood vessels and microglia. We generated endothelial cell (*Cdh5Cre^ERT^*^2^ and *Tie2^Cre^*) ^39, 40^ and vascular mural cell (*Pdgfrb^Cre^*)^41^ *Tgfb1* conditional mutants, and compared these to *Tgfb1^−/−^* mutants as a positive control (Figures 5A and 5B). As previously documented^42^, *Tgfb1^−/−^* mice showed prominent vascular changes (glomeruloid malformations, dilated and tortuous vessels) and associated hemorrhage throughout the cerebrum (asterisk in Figure 5B). Concomitant with vasculopathy and hemorrhage, microglia in *Tgfb1^−/−^*embryos completely lacked P2RY12 staining, even in microglia distant from hemorrhaging (Figure 5B). In comparison,*Tgfb1^fl/fl^;Cdh5^CreER^* endothelial mutants (Figure 5C) and *Tgfb1^fl/fl^;Pdgfrb^Cre^* mural cell mutants (Figures 5D and 5E), both retained P2RY12 expression in microglia, while *Tgfb1^fl/fl^;Tie2-Cre* mutants showed an almost complete loss of P2RY12 expression in these cells (Figures 5F and 5G). We suspect that reduced P2RY12 expression in *Tie2^Cre^* mutants is due to early developmental recombination in, and *Tgfb1* deletion from, microglia precursors; *Tie2^Cre^* is well-known to recombine yolk sac hemogenic endothelium from which microglia are derived.

**Figure 5.**
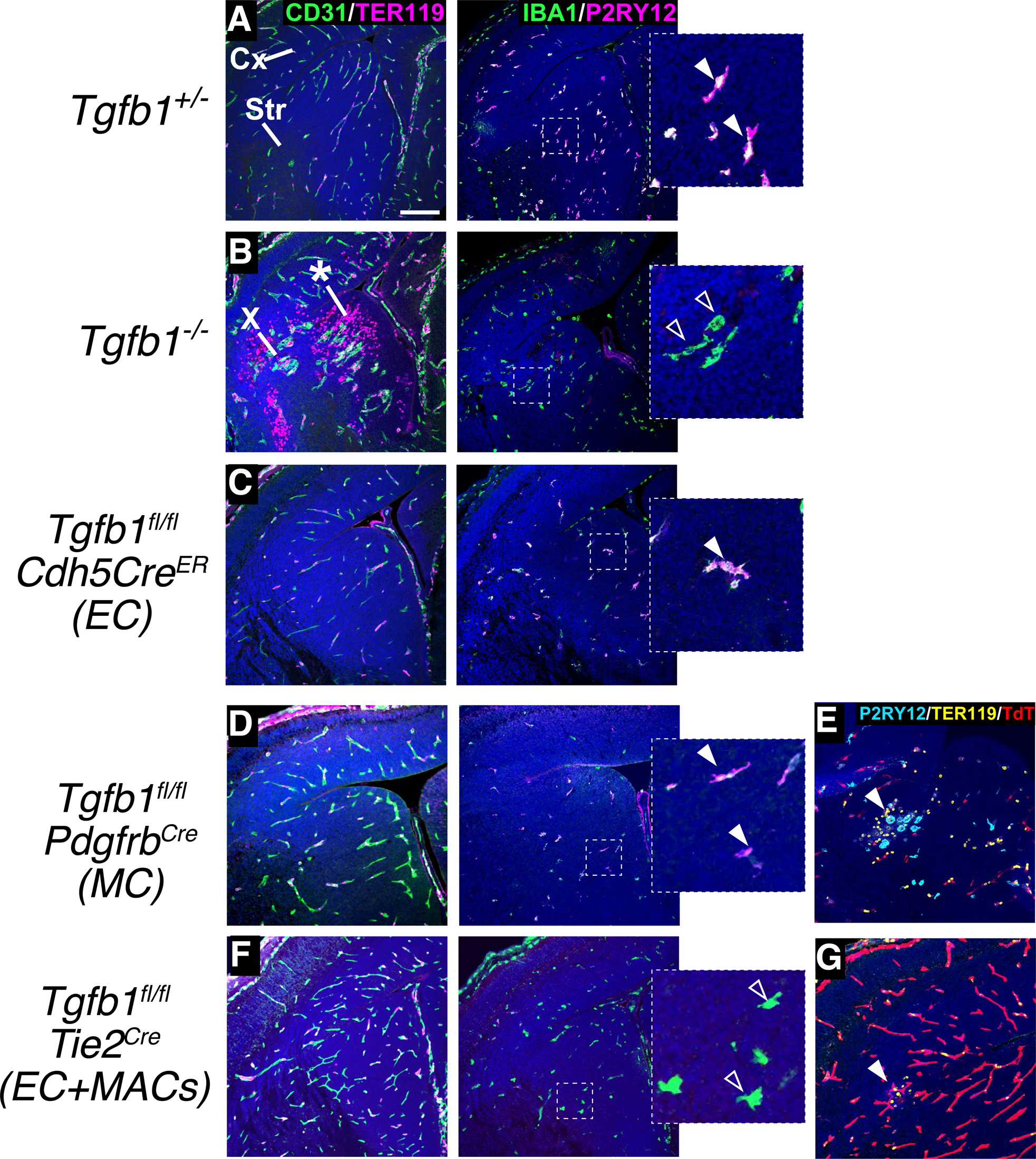
Vascular *Tgfb1* is not required for microglial development. E14.5 coronal brain sections from (**A)** control (*Tgfb^+/−^*) embryos, **(B)** embryos with global (*Tgfb1^−/−^*) or cell-lineage specific deletion of *Tgfb1* (*Tgfb1^fl/fl^*) in **(C)** endothelial cells (*Cdh5Cre^ER^)* **(D,E)** vascular mural cells (*Pdgfrb^Cre^*), or **(F,G)** endothelial cells and microglia/macrophages (*Tie2^Cre^*). Sections were stained for hemorrhage (TER119, magenta) and vasculature (CD31, green) or for committed/hemostatic microglia (IBA1, magenta and P2RY12, green), or to study P2RY12 expression in the context of hemorrhage (P2RY12, cyan and TER119, yellow). Only *Tgfb1^−/−^*mutants have consistent evidence of vascular dysplasia (marked by X) and hemorrhage (asterisk), whereas mice with microglia/macrophage deletion of *Tgfb1* (*Tgfb^−/−^*, and *Tgfb1^fl/fl^*;*Tie2Cre* mutants) have presence of dysmature microglia (open arrowheads, blowups to right). Panels in **E)** and **G)** show *ROSA-TdT (Ai14)* recombination pattern of *Pdgfrb^Cre^* and *Tie2Cre* mouse lines respectively, and P2RY12 expression (or lack thereof) in the sporadic hemorrhage (arrowheads) seen when *Tgfb1* is deleted with these lines. Cx=Cortex, Str=Striatum. Scale bar in A=150*μ*m.

Surprisingly, *Tie2^Cre^*, *Cdh5^CreER^* and *Pdgfrb^Cre^* mutants had few vascular changes (Figures 5C-5F), although we did consistently observe one or two sporadic, small hemorrhages in most *Tgfb1^fl/fl^;Pdgfrb^Cre^*and *Tgfb1^fl/fl^;Tie2^Cre^*embryos (Fig 5E and 5G). Interestingly, in areas of hemorrhage in *Tgfb1^fl/fl^;Pdgfrb^Cre^* embryos, we observed no reduction in P2RY12 expression (Figure 5E), suggesting that hemorrhage and the loss of microglial homeostasis is separable. This observation supports the model that the microglial phenotypes observed in *Itgb8* and *Tgfb1* mutant mice are due to reduced TGFβ1 activation and signaling in microglia, and not secondary consequences of hemorrhage.

### *Tgfb1* is required to maintain postnatal microglial homeostasis

Bulk RNA-seq analysis indicates that *Tgfb1* expression in the adult mouse brain is found primarily in microglia and endothelial cells (Figure 6A)^35^. To assess whether there is an ongoing requirement for *Tgfb1* in postnatal microglial homeostasis we induced recombination in *Tgfb1^fl/fl^;Cx3cr1^CreER^* (Figures 6B-E) and *Tgfb1^fl/GFP^;Cx3cr1^CreER^*(not shown) neonatal mice at P4,5,6. These mice were then collected at P30 for histological analysis. In these neonatally recombined mice we found large patches of dysmature microglia that were characterized by the loss of *Tgfb1* expression and reduced TGFβ signaling (reduced pSMAD3 staining), reactive morphology, loss of homeostatic markers TMEM119 and P2RY12, and upregulation of CD206 (Figures 6B-E). The size and distribution of patches was highly variable, with intermixing of *Tgfb1*-intact and dysmature microglia, suggesting that *Tgfb1*-intact microglia cannot compensate for the loss of *Tgfb1* in adjacent dysmature cells. Similarly, *Tgfb1*-expressing vascular cells were often found intermingled with *Tgfb1*-deleted microglia indicating that *Tgfb1* from endothelial cells cannot compensate for the loss of *Tgfb1* in neighboring cells. Of note, *Tgfb1* tamoxifen-inducible mutants exhibited no obvious neurobehavioral abnormalities, possibly due to variable penetrance and/or an attenuated dysmature microglial phenotypes.

**Figure 6.**
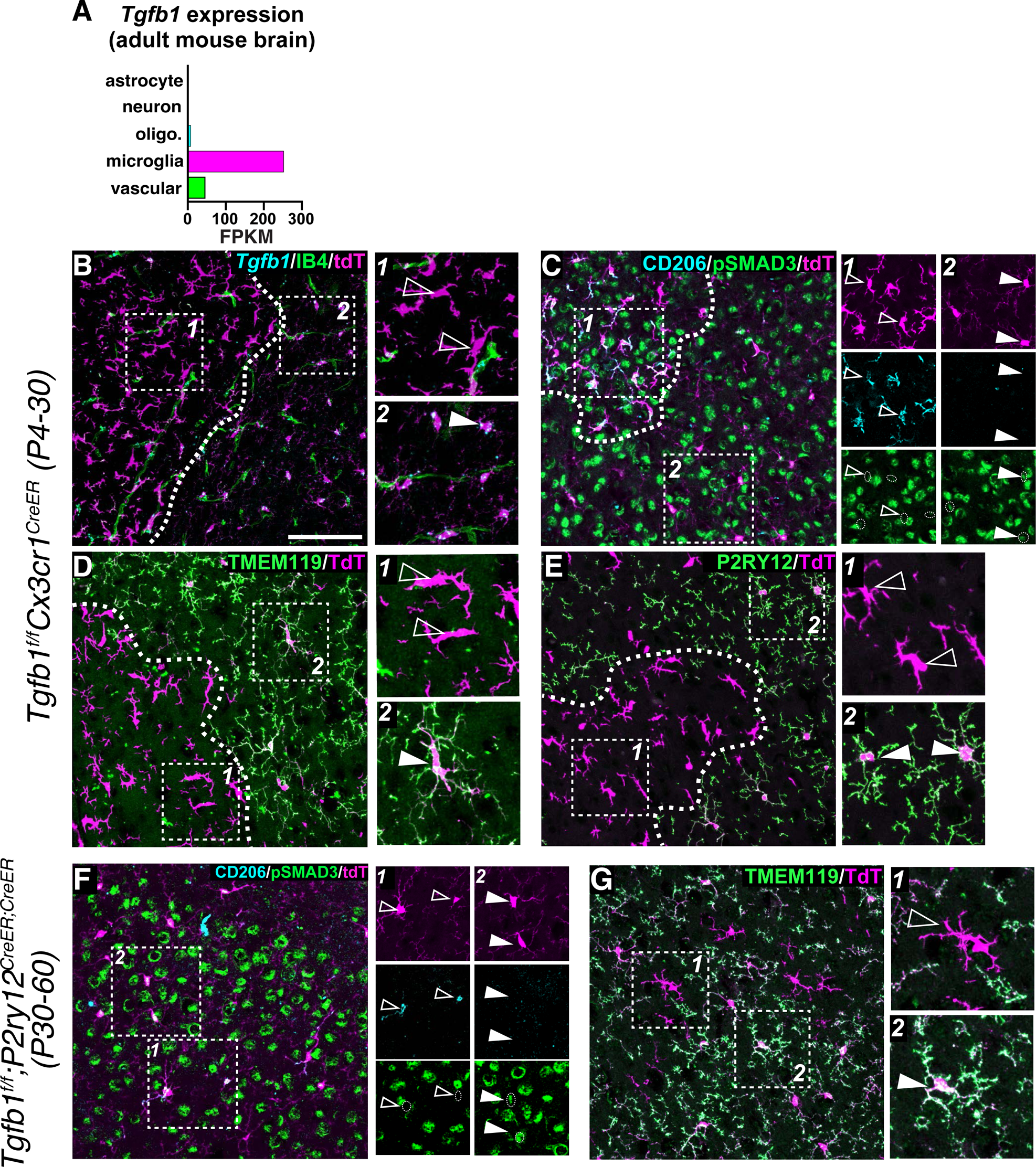
TGFβ1 is required postnatally for microglial homeostasis. **F)** Bulk-seq analysis of *Tgfb1* expression in the adult mouse brain^35^. Analysis revealed enrichment for *Tgfb1* expression primarily in microglia and vascular cells. **B-E)** Analysis of conditional *Cx3cr1^CreER^* mediated deletion of *Tgfb1* deletion in the P30 mouse brain following neonatal tamoxifen induction at P4,5 and 6. Analysis revealed a “patchy” distribution of dysmature microglia characterized by altered morphology and **A)** *Tgfb1* loss; **B)** Upregulation of CD206 and loss of pSmad3 staining; **D)** Loss of the homeostatic marker TMEM119 and **E)** loss of the homeostatic marker P2RY12. Open arrowheads in B-G mark dysmature microglia, closed arrowheads mark dysmature microglia. Postnatal deletion of *Tgfb1* in adulthood (P30-P60) using *P2ry12^CreER^* resulted in isolated production of microglia with altered morphology **(F)**, and loss of pSMAD3 staining **(F)** and downregulation of the homeostatic marker TMEM119 **(G)**, upregulation of CD206 **(H)** Scale bar in B=100*μ*m.

To assess the role of *Tgfb1* in adult microglial homeostasis, we performed experiments to conditionally delete *Tgfb1* in adulthood. Following tamoxifen inductions at P30, we were unable to successfully recombine the *Tgfb1* floxed allele with *Cx3cr1^CreER^*, which we attributed to the large distance between the loxP sites in the *Tgfb1* floxed allele (4.4kb). However, in mice homozygous for *P2ry12-CreER* (*Tgfb1^fl/fl^;P2ry12^CreER/CreER^*and *Tgfb1^f/GFP^;P2ry12^CreER/CreER^*) that were induced starting at P30 and analyzed at P60, we were able to find scattered mosaicly recombined microglia that displayed a similar phenotype to neonatally induced *Tgfb1^fl/fl^;Cx3cr1^CreER^* and *Tgfb1^fl/GFP^;Cx3cr1^CreER^* mice with patchy recombination (reactive morphology, loss of homeostatic markers, upregulation of CD206, and loss of pSMAD3 staining) (Figures 6F, 6G and data not shown). Taken together, these data indicate that microglia require self-produced TGFβ1 to maintain homeostasis.

### Conditional *Smad2/3* mutants reveal role for non-canonical TGFβ signaling in microglial homeostasis

After engaging the TGFβ receptor, TGFβ signaling proceeds through both canonical (SMAD-dependent) and non-canonical (SMAD-independent) pathways (Figure 7A). To understand whether microglia abnormalities and associated neuromotor dysfunction can be attributed to canonical versus non-canonical TGFβ signaling, we generated *Smad2/3^fl/fl^;Cx3cr1^Cre^* mice and compared the behavioral, transcriptional and histological properties of these mice to *Tgfb1^fl/fl^;Cx3cr1^Cre^* mice^43,44^. Neither *Smad2/3^fl/fl^;Cx3cr1^Cre^*or *Tgfb1^fl/fl^;Cx3cr1^Cre^* mutants showed evidence of neonatal brain hemorrhage (data not shown). Following mice into adulthood, *Tgfb1^fl/fl^;Cx3cr1^Cre^* mice presented at 2 months of age with neuromotor dysfunction highly similar to 2-month-old *Itgb8^fl/fl^;Nestin^Cre^*, *Tgfbr2^fl/fl^;Cx3cr1^Cre^* ^7^ and *Lrrc33^−/−^* mice^5^. In contrast, *Smad2/3^fl/fl^;Cx3cr1^Cre^* mice showed no obvious behavioral abnormalities until 15 months of age, at which point some mice exhibited a slight tremor (Figure 7B). None of the *Smad2/3* mutants of the allelic series (e.g. double heterozygotes, homozygous floxed mutant of one floxed *Smad* allele with heterozygosity of the other) showed any behavioral or histological phenotype, suggesting that *Smad2* and *Smad3* can compensate for each other.

**Figure 7.**
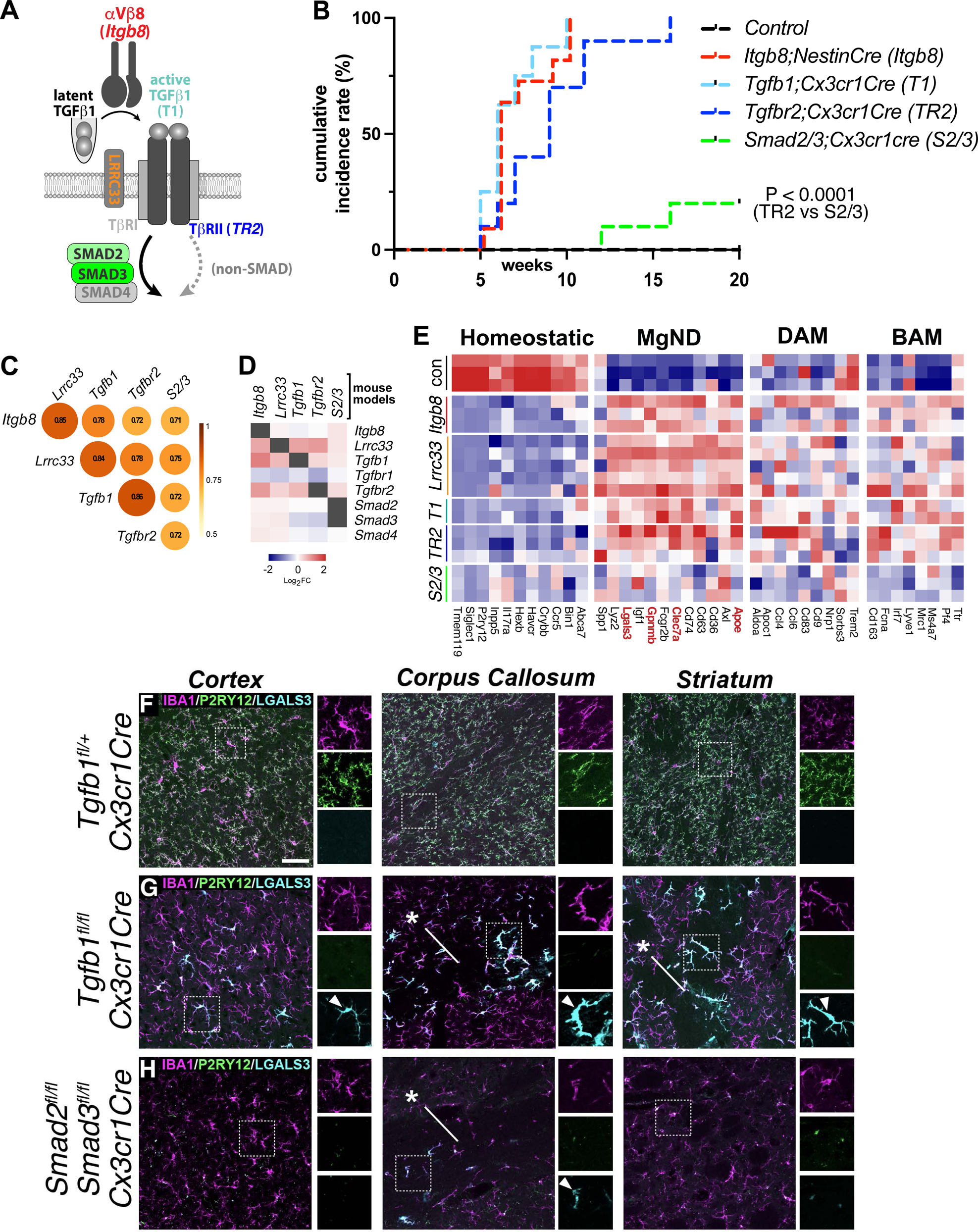
Disruption of non-canonical TGFβ signaling in microglia drives disease-associated gene expression. **A)** Schematic of TGFβ, with mutant mouse models analyzed by bulk and microglial flow cytometry noted by color (*Itgb8*=red; *Tgfb1*=cyan; *Lrrc33*=orange; *Tgfb2*=blue, *Smad2/3*=green). **B)** Cumulative incidence of motor dysfunction seen in different conditional TGFβ pathway mutants as a function of age. Incidence included any of detriment to gait, appearance, or tremor, based on a 0,1,2 rating scale (see ^56^ for details). **C)** Correlation of bulk-seq gene expression across TGFβ mutant models. **D)** Compensatory transcriptional changes of key TGFβ signaling genes in different TGFβ mutant models. **E)** Bulk-seq analysis of microglial homeostatic and disease associated (MGnD/DAM) microglial markers across TGFβ mutant models. **F-G)** Comparison of control **(F)** *Tgfb1^fl+l^;Cx3cr1^Cre^,* **(G)** *Tgfb1^fl/fl^;Cx3cr1^Cre^* and **(H)** *Smad2/3^fl/fl^;Cx3cr1^Cre^* adult mice. Analysis revealed loss of the homeostatic marker P2RY12 in both conditional *Tgfb1* and *Smad2/3* mutants, but significantly higher upregulation of the MGnD-associated microglial marker LGALS3 (see arrowheads). LGALS3 upregulation in *Tgfb1* conditional mutants was significantly higher in white matter (asterisks in **G and H**) and was only seen in the white matter of *Smad2/3* conditional mutants (**H**). Scale bar in F=50*μ*m.

To understand why *Smad2/3* double mutants lacked neurobehavioral symptoms comparable to other *Itgb8-Tgfb1* signaling pathway mutants, we compared the transcriptional properties of *Itgb8*, *Lrrc33*, *Tgfb1*, *Tgfbr2* and *Smad2/3* conditional mutant models using whole brain bulk RNASeq (Figures 7C-E, S5, S6 and S7). Whole brain transcriptomes from TGFβ pathway mutants were highly correlated, as would be expected from disrupting genes in a shared signaling pathway (Figure 7C). Interestingly, *Lrrc33* and *Itgb8* mutants, the two most “proximal” to TGFβ1 activation, were the two most highly correlated gene sets. In contrast, *Smad2/3* mutants, being the most distal from *Tgfb1* activation were the least correlated, possibly reflecting the accumulation of interceding non-canonical signaling pathways. We also identified several key differences among pathway mutants. First, we noticed that the pathway genes themselves were differentially altered in the various mutants (Figure 7D) suggesting compensatory pathway feedback; this compensation was lowest in the *Smad2/3* mutants. Beyond these potential feedback-associated gene changes, the most highly differentially expressed genes were largely microglia-specific, which we confirmed by comparing to bulk RNAseq of sorted microglia from each model (Figures 7E, S5). Notably, while the reduction in homeostatic genes in *Smad2/3* mutant mice was comparable to other models, increases in MGnD/DAM and BAM genes were generally less pronounced in *Smad2/3* mutants than in all other mutants (Figure 7E). Genes associated with astrocytosis were also not as highly upregulated in *Smad2/3* mutants as compared to other TGFβ mutant models (*ApoE*, *GFAP*, *Fabp7*, and *Vimectin)* in our bulk RNA-seq analysis (Figure S6). Focusing on genes that were differentially expressed in *Tgfb1-* and *Tgfbr2-*, but not *Smad2/3*-mutants yielded several putative biological processes enriched in symptomatic mutants. Myelination and complement associated synaptic pruning were among the most representative terms associated with *Tgfb1^fl/fl^;Cx3cr1^Cre^*and *Tgfbr2^fl/fl^;Cx3cr1^Cre^* microglia and whole brain RNAseq gene sets, which also differentiate these models from *Smad2/3^fl/fl^;Cx3cr1^Cre^*mice (Figure S5).

Among microglia-specific genes differentially expressed in *Smad2/3^fl/fl^;Cx3cr1^Cre^*versus other mutant models, we found *Apoe*, *Clec7a* and *Lgals3* to be particularly interesting (Figure 7E). While considered canonical markers of DAM/MGnD microglia, these genes are also transiently enriched in populations of developmentally restricted microglia associated with myelin and axonal tracts (Axonal tract microglia, ATM)^45^, and in areas of active neurogenesis (proliferation associated microglia, PAM)^46^. Of note, we previously found evidence for delayed myelination in *Itgb8^fl/fl^;Nestin^Cre^* and *Tgfbr2^fl/fl^;Cx3cr1^Cre^*mutant mice^7^. The expression of these genes in MGnD microglia depend on the APOE pathway: MGnD signature genes including *Spp1*, *Gpnmb,* and *Clec7a* are suppressed in disease models that also lack *Apoe*^28,47^, and expression of human APOE isoforms APOE ε4 in microglia similarly suppresses MGnD polarization relative to the ε3 isoform (Yin et al. Nature Immunology, In Press). In contrast, developmental expression of MGnD/ATM genes near myelin/axonal tracts does *not* depend on *Apoe*^46^.

We were therefore interested to understand the degree to which *Apoe* might regulate PAM/MGnD signature genes, and whether increased expression of APOE in *Itgb8*, *Lrrc33*, *Tgfb1*, and *Tgfbr2* mutants might account for their more severe neuromotor symptoms. To that end, we performed an epistatic knockout analysis by deleting *Tgfbr2* or *Apoe* singly or together. We created *Tgfbr2^fl/fl^;Cx3cr1^Cre^;Apoe^+/−^* and *Tgfbr2^fl/fl^;Cx3cr1^Cre^;Apoe^−/−^* mutant mice, and evaluated their behavioral and microglial phenotypes at P90^48^ (Figure S7). Surprisingly, despite the transcriptomic similarity between dysmature and MGnD microglia, we found that deletion of APOE in *Tgfbr2^fl/fl^;Cx3cr1^Cre^*mice had no major effects on neuromotor or microglial phenotypes (Figure S7). This is similar to PAM/ATM microglia which also do not depend on APOE^46^. These data indicate that, unlike MGnD microglia, dysmature microglia do not depend on APOE for their polarization to occur, or alternatively, that non-canonical TGFβ signaling functions directly downstream of *Apoe* in microglial polarization.

Compared to TGFβ-pathway mutants, we observed a similar pattern of polarization among most MGnD and BAM markers, suggesting that phagocytic microglia, especially those carrying the APOE ε3 allele, transition to a BAM-like state, whereas microglia carrying the APOE ε4 allele are less BAM-like (Figure S8A). As microglia lacking TGFβ signaling adopt a similar BAM-like state, we analyzed microglial RNA expression data following a Aβ plaque-reducing treatment of ITGB8-blocking antibody in an Alzheimer’s disease (AD) murine model (APP/PS1 mice), also from our recent study (Yin et al. Nature Immunology, In Press). Interestingly, we observed upregulation of some BAM markers after ITGB8 block, although to a lesser extent than genetic inactivation of components of the TGFβ cascade (Figures 7 and S8B). We previously showed that blocking ITGB8 in the brain induced the expression of genes involved in antigen processing and interferon-γ in microglia (Yin et al. Nature Immunology, In Press) (Figure S8C). This pattern was also observed in mutant microglia lacking proximal components of TGFβ signaling (Figure S8D), although it was much lower in *Smad2/3* mutants, suggesting that antigen processing and interferon-γ signaling could be relevant for the neuromotor phenotype of fulminant TGFβ mutant models.

To determine the underlying causes of the motor dysfunction seen in TGFβ pathway mutants, we considered the non-autonomous roles of molecules secreted by dysmature microglia. In AD mouse models, astrocyte activation is correlated with the degree of MGnD microglial polarization, and is mechanistically linked to microglia expression of LGALS3 (Yin et al. Nature Immunology, In Press). Examination of LGALS3 expression by immunohistochemistry confirmed that LGALS3 expression was much higher in *Tgfb1^fl/fl^;Cx3cr1^Cre^* than *Smad2/3^fl/fl^;Cx3cr1^Cre^*mice, providing a potential explanation for different behavioral outcomes in these mice, despite the loss of microglial homeostatic markers in both models (Figures 7F-H), and in line with our observations regarding astrocytosis-related gene expression (Figure S6). Interestingly, while most brain regions in *Smad2/3^fl/fl^;Cx3cr1^Cre^* mice did not show microglial LGALS3 expression, areas containing dense white matter, such as the corpus callosum, showed upregulation of LGALS3 (asterisk in Figure 7H). *Tgfb1^fl/fl^;Cx3cr1^Cre^* also had much higher relative levels of microglial LGALS3 expression (arrowheads in Figure 7) in the corpus callosum and fiber tracts of the striatum (arrowheads in Figure 7G), suggesting that white matter microglia may be particularly reactive to the effects of TGFβ signaling disruption. Together, these data indicate that isolated disruption of canonical (SMAD-directed) TGFβ signaling in microglia results in neuropathologial phenotypes that are less severe than in mice with more proximal disruptions in the TGFβ signaling cascade, and that the disease-associated microglial gene LGALS3 may drive astrocytic changes seen in *Tgfb1^fl/fl^;Cx3cr1^Cre^*mice that are less severe or absent in *Smad2/3^fl/fl^;Cx3cr1^Cre^*mice.

## Discussion

Here, we show that that microglial differentiation is dependent upon the activation of microglial TGFβ1 by *Itgb8*-expressing radial glia progenitors. Disruption of this signaling by conditionally deleting radial glia progenitor *Itgb8*, microglial *Tgfb1*, or the downstream transcription factors *Smad2/3* in microglia results in a microglial transcriptional phenotype characterized by the absence of mature microglial markers and persistent expression of genes normally expressed in microglial precursors and immature microglia.

Interestingly, we find that domain-specific deletion of *Itgb8* in early embryonic radial glia progenitors results in microglial defects that are restricted to the brain regions created by these progenitors. Furthermore, consistent with recent reports that microglia are derived from a CD206+ precursor^22–24^ and the earliest characterizations of microgliogenesis by del Rio-Hortega^25^, we find that embryonic radial glial endfeet make direct contact with CD206+ meningeal macrophages. Together, our data suggest that the physical interaction between radial progenitors and immature microglia is crucial for radial glial *Itgb8*-mediated TGF-β signaling in microglia. This is consistent with previous reports of microglia/radial glia interactions^49^. The lack of a similar microglial phenotype in *Itgb8^fl/fl^;hGFAP^Cre^*mutants, which recombine dorso-lateral radial progenitors ∼3 and ∼5 days after *Nestin^Cre^* and *Emx1^Cre^* respectively, places the timing of this interaction between E9 and E13.5, when microglia initially begin to invade the developing nervous system^34,14^. Remarkably, dysmature microglia in *Itgb8/Tgfb1* pathway mutants are quite similar to myeloid progenitors at this age. Our data support the model that microglia are derived from a CD206+ precursor and that disruption of *Itgb8/Tgfb1* signaling results in the blockage of microglial marker expression and the maintained expression of immature markers that would otherwise be downregulated. Our analysis shows that this is reflected by widespread epigenetic changes to homeostatic, MGnD/DAM and BAM gene loci. These epigenetic changes are long-lasting and stable through adulthood, beyond the point at which *Itgb8* appears to be necessary for microglial homeostasis. This suggests that the epigenetic changes in *Itgb8^fl/fl^; Emx1^Cre^*and *Itgb8^fl/fl^; Nestin^Cre^* mice may be irreversible, although this remains unclear and is an area for future investigation. Additional future studies are needed to determine where and more precisely when microglial/radial glia interaction occurs developmentally, and also whether the specific interactions between ingressing microglia and radial glial basal and/or apical endfeet in the meninges or ventricular surface are necessary for microglial differentiation.

Our findings suggest that additional new models of brain region restricted microglial dysfunction can be created by deleting *Itgb8* in other radial progenitor domains. Taking advantage of the integrin α_V_β_8_ trans-activation of TGFβ1 to create genetic models of brain region-restricted microgliopathy is particularly useful, as microglial *Cre* and *CreER* lines generally recombine microglia throughout the nervous system, and not in particular brain regions^37,50,51,52^. We believe that the study of brain region-restricted microglial dysfunction will provide crucial information about how dysmature microglia in particular brain regions contribute in a modular fashion to more complex behavioral and cognitive disruptions seen in other animal models and in humans with pervasive neurodevelopmental immune dysfunction. Combining mouse models of neuroinflammatory or neurodegenerative disease with brain region-specific *Itgb8* conditional knock-out models will be especially illuminating.

In line with recent cryoEM data^10,11^, we show that microglia depend on cell-autonomous expression of *Tgfb1* for their early maturation. Interestingly, we only observe a fulminant vascular/hemorrhage phenotype in *Tgfb1* null mice; deletion of *Tgfb1* in endothelial cells and microglia (*Tie2^Cre^*), or in vascular mural cells and fibroblasts (*Pdgfrb^Cre^*) had no major effect on cerebrovascular morphogenesis or barrier function. Therefore, vessel-expressed TGFβ1 is apparently dispensable for microglial maturation, but the source of TGFβ1 for vascular maturation remains unknown. Fitting with the paracrine activation / autocrine signaling model, we propose that radial glia ITGB8 activates TGFβ1 on endothelial cells and/or pericytes, so that the loss of endothelial *Tgfb1* can be compensated by pericyte TGFβ1 or vice versa. It is also possible that TGFβ1 in the embryonic circulation can compensate for the loss of vascular TGFβ1^42^.

In contrast to *Itgb8*, which we show is largely dispensable for maintaining microglial homeostasis, we find that microglial *Tgfb1* is necessary for microglial homeostasis postnatally. One potential explanation for this discrepancy could be the activation of TGFβ1 by other integrin complexes or by integrin-independent pathways postnatally. We recently found that antibody-based blockade of ITGB8 in AD mouse models enhances MGnD polarization and reduces the size of amyloid plaques (Yin et al. Nature Immunology, In Press). Interestingly, conditional deletion of *Lrrc33* from microglia/macrophages using *Cx3cr1^CreER^* resulted in no major transcriptional or behavioral phenotypes compared to mice with global or conditional deletion of *Lrrc33* during development, which virtually phenocopy *Itgb8^fl/fl^; Emx1^Cre^* and *Itgb8^fl/fl^; Nestin^Cre^* mutants^5^. This raises the possibility that proteins that present/modify *Tgfb1* are only active developmentally or in disease contexts, an important consideration for the design of TGFβ-targeting therapeutics.

In line with previous reports^2,30,45,53^, our study emphasizes the common features of developmental and disease-associated microglia. In particular, by comparing *Smad2/3* conditional mutants to other conditional TGFβ mutant models, we find that there is a phenotypic gradient characterized by lower vs higher relative expression of MGnD and BAM genes and directly correlated neuromotor dysfunction. We propose that polarization along this gradient is regulated by non-canonical TGFβ signaling, a model that is supported by previous observations that disruption of non-canonical TGFβ signaling promotes microglial homeostasis^54,55^. Importantly, while the dysmature gene signature may be protective in neurodegenerative disease, when it is present during development, mice develop severe neurological impairments. Additional studies are needed to better understand the nuance of how microglial reactivity is regulated in different brain regions during development and how these same gene regulatory pathways manifest in the context of brain injury and disease.

## Acknowledgements

This study was in supported in part by HDFCC Laboratory for Cell Analysis Shared Resource Facility through a grant from NIH (P30CA82103). This research was supported, in part, by NIH grants R01NS119615, R01NS123168 (T.D.A.) and the UCSF Vision Core shared resource of the NIH/NEI P30 EY002162. We would like to thank Marie La Russa for helpful manuscript editing suggestions. We would like to thank Dr. Richard Flavell for providing *Tgfb1^flox^* and *Tgfb1^GFP^* mice. We would also like to thank Dr. Rosemary Ackhurst for providing *Tgfb1* null mice. Microglial cartoons in the Graphical Abstract were created with BioRender.com.

## Author Contributions

T.A. supervised the study, performed the behavioral analysis in Figure S1 and did the cell counting in Figure S2. G.L.M. designed the study, under the supervision of T.A. G.L.M. performed the histological analysis, with specific contributions from others listed here. N.S. performed preliminary histological analysis of postnatal *Tgfb1^fl/fl^/Tgfb1^F/GFP^; Cx3cr1^Cre^* and *Smad2/3^fl/fl^; Cx3cr1^Cre^* mice. G.L.M performed the imaging, with support from T.A. for Figure S2. Mouse husbandry was done by G.L.M., with support from N.S., who initially bred *Tgfb1^fl/fl^/Tgfb1^F/GFP^; Cx3cr1^Cre^* and *Smad2/3^fl/fl^; Cx3cr1^Cre^* mice, and with support from A.K., L.T., M.B., K.C, and D.M.. G.L.M. performed RNA isolation and bulk RNA-seq experiments for Figure 7. N.S. performed bioinformatic analysis in Figure 2 and 7. X.Z. performed RNA-seq, ATAC-seq and ChIP-seq analysis of sorted microglia in *Itgb8^fl/fl^;Emx1^Cre^*mice for Figures 2 and 3, with bioinformatic support from K.K. A.K and L.T sectioned *Tie2-Cre* and *PdgfRb-Cre* embryos used in Fig 3. G.L.M. performed the flow cytometry experiments to isolate microglia from *Tgfb1^fl/fl^; Cx3cr1^Cre^* mice, with support from C.O. M.L. created the graphical abstract and helped to edit the manuscript. C.O. performed the flow cytometry experiments to isolate microglia from *Smad2/3^fl/fl^; Cx3cr1^Cre^* mice. L.T. collected and stained embryos used in the *Emx1-Cre; RG-Brainbow* analysis. H.K. helped to collect *Nestin^Cre^* embryos used in Fig 1. H.P. provided the *Itgb8-TdT* mouse line and E.S. provided *RG-Brainbow* mouse line. K.A. supervised C.O. D.S. provided input regarding experimental design and interpretation. O.B. supervised work by X.Z. and K.K. The manuscript was written by G.L.M and T.A.

## Methods

### STAR Methods Table

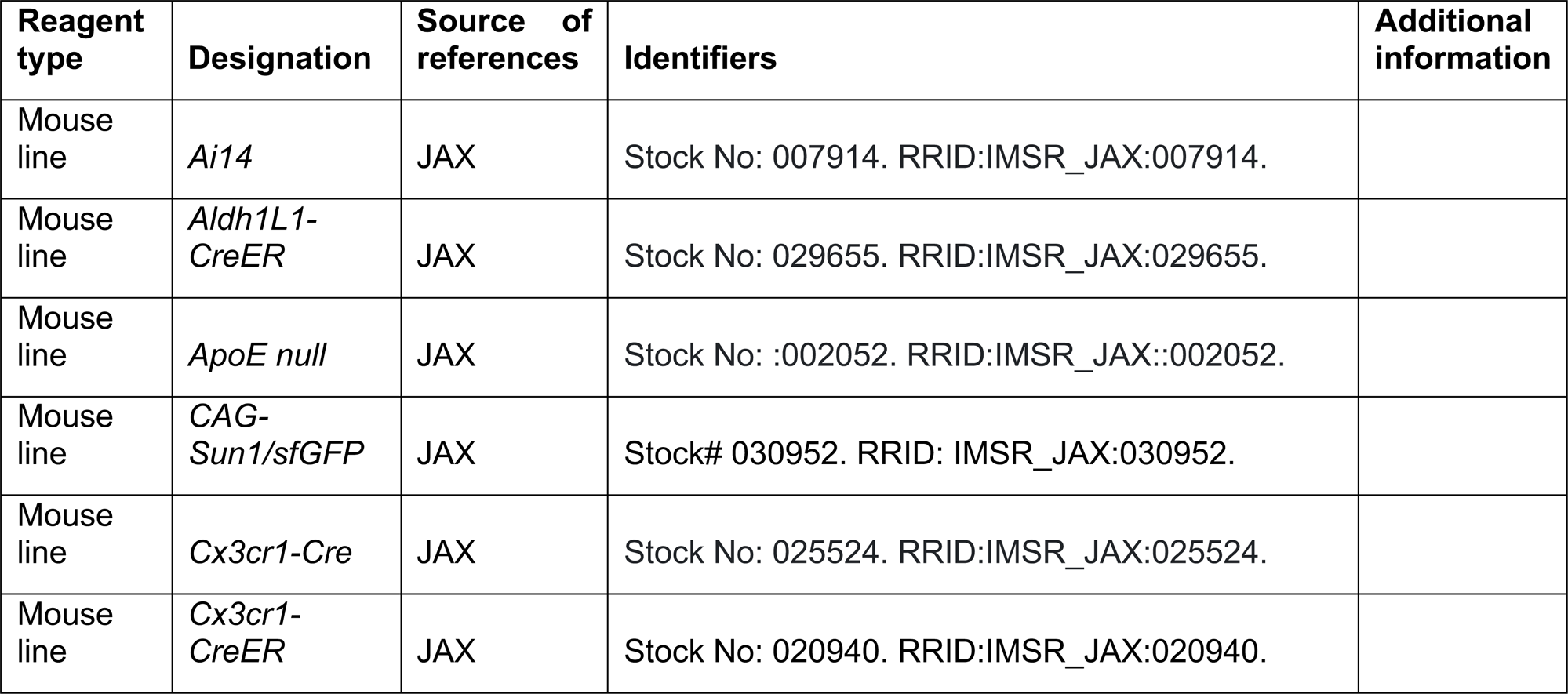
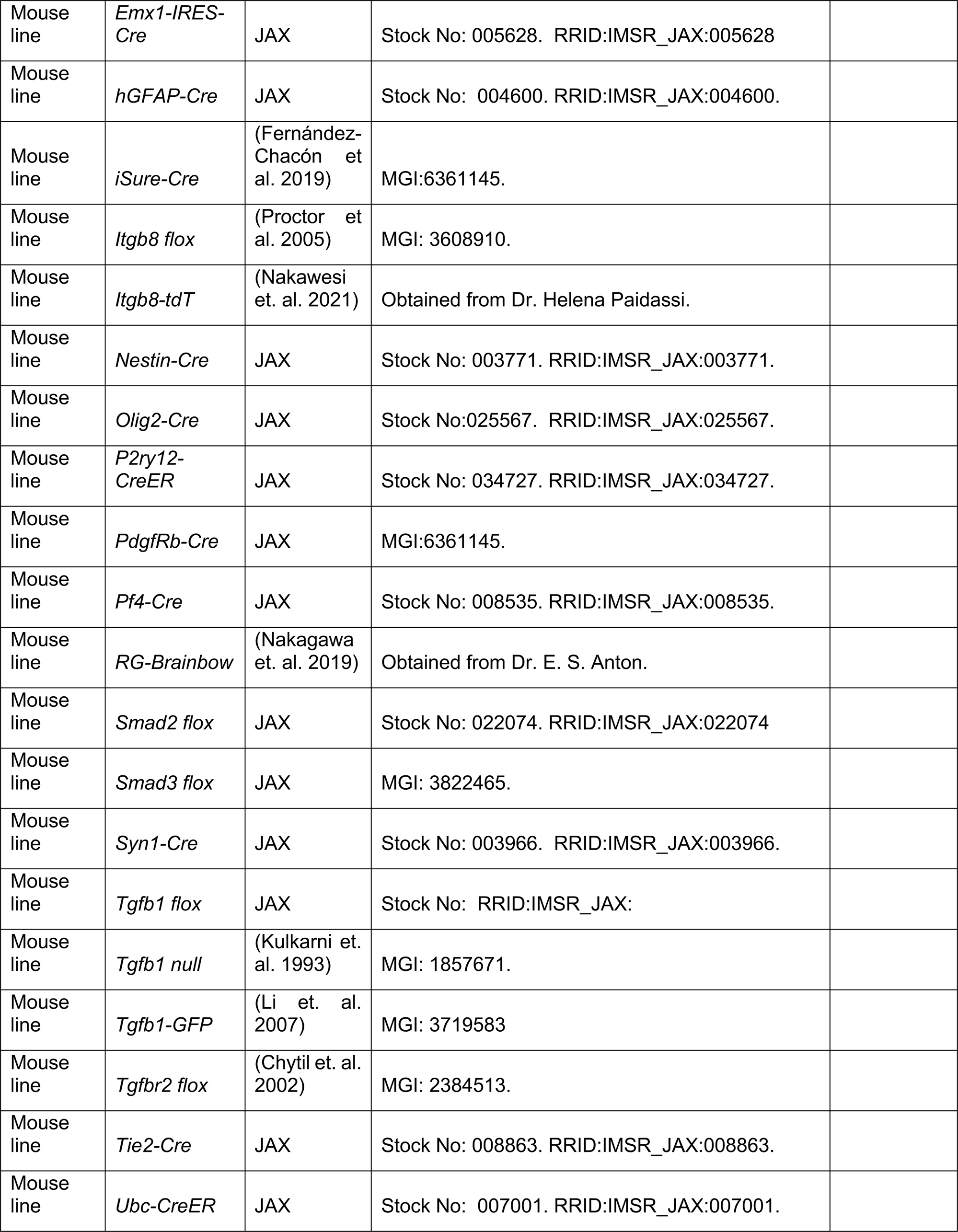
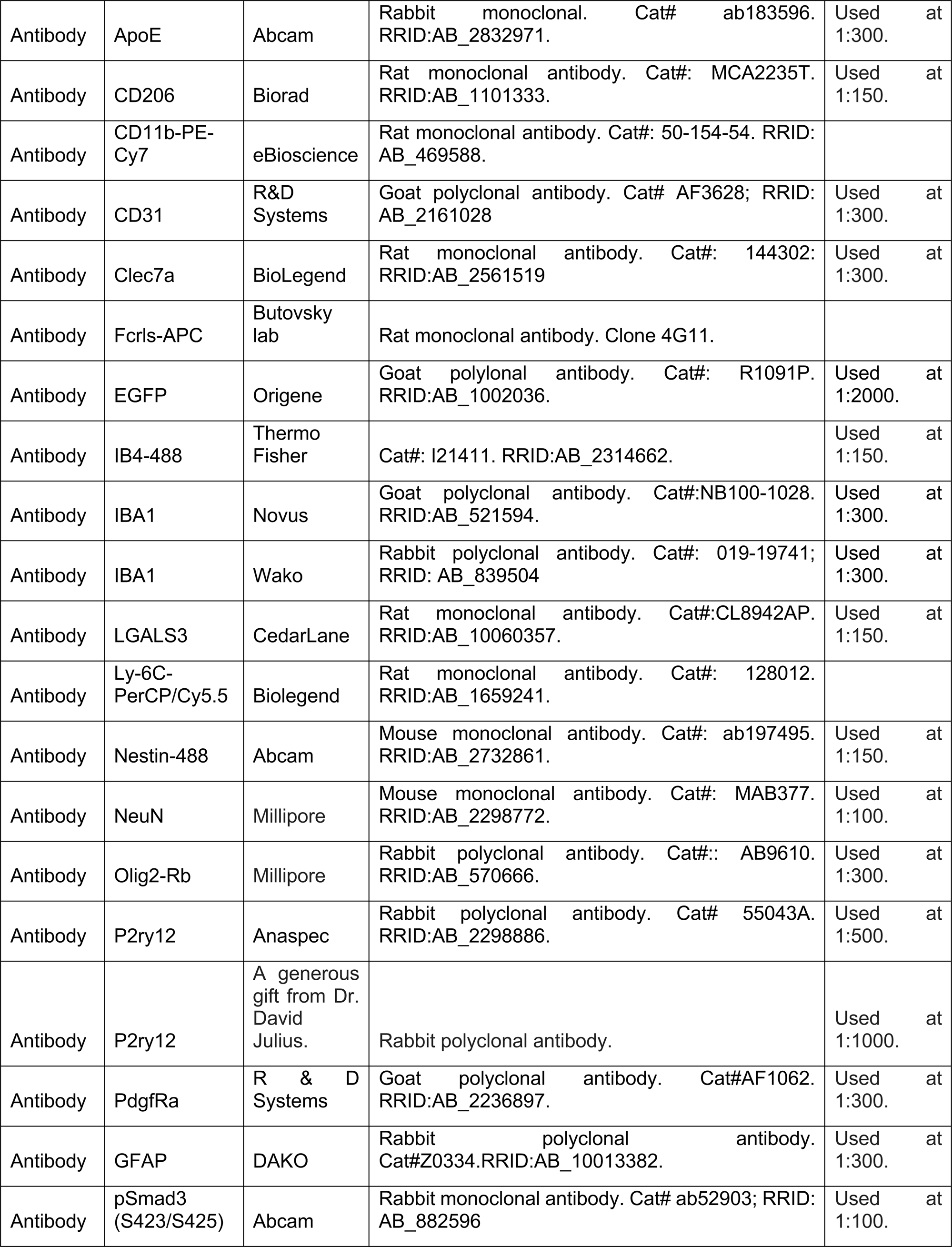
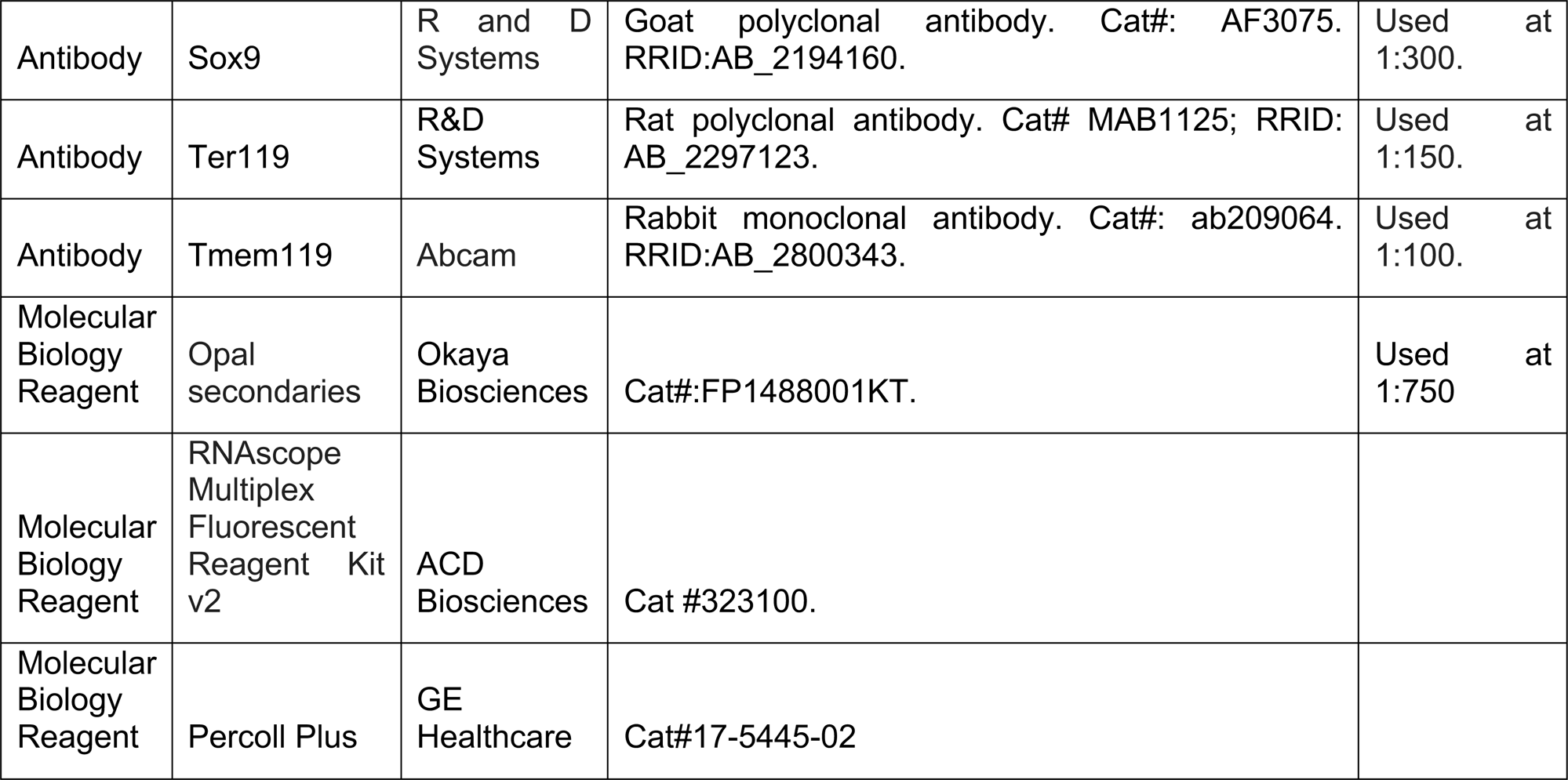

### Mice

All mouse work was performed in accordance with UCSF Institutional Animal Care and Use Committee protocols. Please see key resources table for additional mouse line information. All TdT reporter analysis was done with the *ROSA-TdT* (*Ai14)* mouse line, except for the embryonic *Nestin^Cre^*mice in Figure 1 and *Syn1^Cre^* mice in Figure S4, which were crossed to the membranous TdT reporter expressing Cre-dependent *iSuRe-Cre* mouse line.

### Histology and Immunostaining

Adult mouse mouse brains were harvested at P60-P90 following transcardial perfusion with 20 mL cold PBS and 20 mL cold 4% formaldehyde. Brains and embryos were fixed overnight at 4 degrees in 4% formaldehyde, followed by overnight incubation in 30% sucrose. Samples were embedded (Tissue Plus O.C.T. Compound, Fisher Scientific) and sectioned at 20 µm. Some adult brains were sectioned at 40um. Sections were immunostained using a blocking/permeabilization buffer of PBS containing 2% BSA, 5% donkey serum and. 5% TritonX-100. Primary and secondary antibodies were diluted in PBS containing 1% BSA and. 25% TritonX-100. Secondary antibodies conjugated to Alexa fluorophores were used at 1:300 (Jackson ImmunoResearch). Immunostained sections were mounted with Prolong Gold Antifade Mountant (Thermofisher P36930). Please see STAR Resources Table for list of antibodies and the relevant concentrations used for immunofluorescence.

### RNAscope

20um cryosectioned tissue sections were processed for RNAscope using the manufactures protocol for cryosectioned tissue and reagents from the RNAscope Multiplex Fluorescent Reagent Kit v2 (Cat #323100) and Opal secondaries (Okaya Biosciences FP1488001KT). A custom RNAscope probe set was used to target the floxed exon 1 of *Tgfb1* (Cat #1207831-C1). Slides were hybridized in the ACD HybEZ II Oven (Cat #321711).

### Tamoxifen induction

Recombination was induced by three doses of tamoxifen dissolved in corn oil, administered by oral gavage every other day (150 µL of 20 mg/mL). For embryonic mouse inductions, pregnant dams were given tamoxifen (150 µL of 20 mg/mL) on E11.5, E12.5 and E13.5, for a total of three gavage injections. Neonatal tamoxifen injections were done at P4,5, and 6, at a dosage of 500*μ*g injected intraperitonealy (50uL of a 10mg/mL solution in corn oil). All mouse work was performed in accordance with UCSF Institutional Animal Care and Use Committee protocols. Mice had food and water ad libitum.

### Flow cytometry

The *Itgb8^fl/fl^;Emx1^Cre^*Mice were euthanized in a CO_2_ chamber and then transcardially perfused with 10 ml cold Hanks’ Balanced Salt Solution (HBSS, ThermoFisher, 14175103). The mouse brain was isolated and the cortex was dissected from one hemisphere for microglia purification using the standardard isolation procedure established in Butovsky lab^57^ Briefly, the cortex was homogenized and resuspended with 5 ml 70% Percoll Plus (GE Healthcare, 17-5445-02) and 5 ml 37% Percoll Plus placed on top. The microglia were enriched in the interface layer after centrifugation in 800 g, 4°C, for 25 min with an acceleration of 2 and a deceleration of 1. The microglia enriched cell population were stained with PE-Cy7 anti-mouse CD11b (1:300, eBioscience, 50-154-54), APC anti-mouse Fcrls (1:1000, clone 4G11, Butovsky Lab), and PerCP/Cy5.5 anti-mouse Ly-6C (1:300, Biolegend, 128012). The cells were then processed by a BD FACSAria^TM^ II (BD Bioscience) and CD11b^+^Fcrls^+^ Ly-6C^−^ cells were sorted into Eppendorf tubes for RNA-seq. Microglia from *Smad2/3^fl/fl^;Cx3cr1^Cre^* and *Tgfb1^fl/fl^;Cx3cr1^Cre^* mice were isolated as previously described by the Arnold lab^51^.

### RNA sequencing

For the sequencing of microglia isolated from *Itgb8^fl/fl^; Emx1^Cre^* mice, one thousand sorted microglia resuspended in 5 ml TCL buffer with 1% 2-Mercaptoethanol were placed into a 96-well plate and sent to Broad Technology Labs for Smart-Seq2 following their standard protocol. Briefly, cDNA libraries were generated using the Smart-seq2 protocol^58^. The amplified and barcoded cDNA libraries were loaded into Illumina NextSeq500 sequencer using a High Output v2 kit to generate 2 × 25 bp reads. For whole brain samples, RNA isolation was performed using dounce homogenization of dissected cerebral cortex samples, followed by Trizol extraction and alcohol precipitation. For Total RNA from FACS-sorted microglia from *Smad2/3^fl/fl^;Cx3cr1^Cre^* and *Tgfb1^fl/fl^;Cx3cr1^Cre^* mice, RNA was isolated using the QIAGEN RNAeasy micro kit. PolyA+ unstranded libraries were synthesized from total RNA (RIN > 6) with NEBNext Ultra II RNA Library Prep Kit and sequenced in a Illumina HiSeq4000 system (150 pb paired-end setting). Demultiplexed fastq files were aligned to the mouse genome (mm10) with Rsubread and quantified with FeatureCounts. Count normalization and differential expression analysis were performed with DESeq2. Correlations between multiple datasets were computed with the cor() function from base R and visualized using the corrplot package. Heatmaps were generated with the pheatmap package. Venn diagrams were produced with standard GNU coreutils and drawn in Inkscape. Overrepresentation analysis was performed using PANTHER from the GO portal. Datasets used in this study are as follows: newly generated data (whole brain data, isolated *Tgfb1* mutant microglia data) have been deposited into the GEO database and will be made public upon publication. *Itgb8^fl/fl^;Emx1^Cre^*microglia and *Smad2/3^fl/fl^;Cx3cr1^Cre^* data were retrieved from the GEO database and will also be made public upon publication. *Tgfbr2^fl/fl^;Cx3cr1^Cre^* microglia is from GEO GSE124868.Data from *Lrrc33−/−* microglia and whole brain was obtained from GEO GSE96938.

For bulk RNA samples, RNA samples were quantified using Qubit 2.0 Fluorometer (ThermoFisher Scientific, Waltham, MA, USA) and RNA integrity was checked using TapeStation (Agilent Technologies, Palo Alto, CA, USA). The RNA sequencing libraries were prepared using the NEBNext Ultra II RNA Library Prep Kit for Illumina using manufacturer’s instructions (New England Biolabs, Ipswich, MA, USA). Briefly, mRNAs were initially enriched with Oligod(T) beads. Enriched mRNAs were fragmented for 15 minutes at 94 °C. First strand and second strand cDNA were subsequently synthesized. cDNA fragments were end repaired and adenylated at 3’ends, and universal adapters were ligated to cDNA fragments, followed by index addition and library enrichment by PCR with limited cycles. The sequencing libraries were validated on the Agilent TapeStation (Agilent Technologies, Palo Alto, CA, USA), and quantified by using Qubit 2.0 Fluorometer (ThermoFisher Scientific, Waltham, MA, USA) as well as by quantitative PCR (KAPA Biosystems, Wilmington, MA, USA).

Ultra-low input RNA sequencing libraries (from sorted *Tgfb1^fl/fl^;Cx3cr1^Cre^*microglia) were prepared by using SMART-Seq HT kit for full-length cDNA synthesis and amplification (Takara, San Jose, CA, USA), and Illumina Nextera XT (Illumina, San Diego, CA, USA) library was used for sequencing library preparation. Briefly, cDNA was fragmented, and adaptor was added using Transposase, followed by limited-cycle PCR to enrich and add index to the cDNA fragments. The sequencing library was validated on the Agilent TapeStation (Agilent Technologies, Palo Alto, CA, USA), and quantified by using Qubit 2.0 Fluorometer (ThermoFisher Scientific, Waltham, MA, USA) as well as by quantitative PCR (KAPA Biosystems, Wilmington, MA, USA). The sequencing libraries for bulk and ultra-low samples were multiplexed and clustered onto a flowcell on the Illumina NovaSeq instrument according to manufacturer’s instructions. The samples were sequenced using a 2×150bp Paired End (PE) configuration. Image analysis and base calling were conducted by the NovaSeq Control Software (NCS). Raw sequence data (.bcl files) generated from Illumina NovaSeq was converted into fastq files and de-multiplexed using Illumina bcl2fastq 2.20 software. One mis-match was allowed for index sequence identification.

### Imaging

Confocal images were taken with 10x, 20x or 40x objectives using a motorized Zeiss 780 upright laser scanning confocal microscope or a motorize Zeiss 900 inverted laser scanning confocal microscope. Image brightness and contrast was optimized in ImageJ. Images and figures were arranged in Adobe Illustrator.

### Statistical analysis

Sample size was not precalculated using a statistical framework, but conformed to general standards in the field. Sample analysis was not blinded. For all immunostaining quantification, values for each mouse were calculated by averaging three pictures from each mouse. Differences between means were compared using a two-tailed t-test, with an alpha of 0.05.

### ATAC-seq

ATAC-seq was carried out following the Omni-ATAC-seq protocol^59^. Briefly, 50k cortical microglia (CD11b^+^Fcrls^+^ Ly-6C^−^) from seven *Itgb8^fl/fl^;Emx1^Cre^* P60-P90 mouse brains and seven control littermates were sorted into Eppendorf tubes by a BD FACSAria^TM^ II (BD Bioscience). Samples from each mouse served as a separate biological and experimental replicate. After being washed with cold dPBS, microglia were permeabilized with ATAC-Resuspension Buffer containing 0.1% NP40, 0.1% Tween-20, and 0.01% Digitonin and washed with ATAC-RSB buffer with 0.1% Tween-20. The isolated microglial nuclei were then incubated in transposition mixture at 37℃ for 60 min. The DNA were purified using a Zymo DNA clean and concentrator-5 kit (Zymo research, Cat.#D4014) and amplified using NEBNext^®^ High-Fidelity 2x PCR Master Mix (NEB, Cat. #M0541s) for 9 cycles. The ATAC libraries were size selected using SPRIselect beads (Beckman Coulter, Cat.#B23317) and quantified using Agilent 2100 Bioanalyzer and Qubit (Invitrogen, Cat.#Q33238). The pooled libraries were then sent to Azenta Life Sciences for paired-end sequencing (2×150 bp) on Illumina HiSeq platform.

### ChIP-seq

Anti H3K9ac ChIP-seq were performed using using the iDeal ChIP-seq kit for Histones (Diagenode, Cat.#C01010059) and NEBNext® UltraTM II DNA Library Prep Kit for Illumina® (NEB, Cat.#E7103S). Briefly, microglia were sorted from *Itgb8^fl/fl^;Emx1^Cre^* mouse brain cortex and were pooled from 3-4 different individuals with same genotype to achieve > 200 k cells per biological replicate. 3 mutant and 4 littermate control (*Emx1-^Cre(–)^*, *Itgb8^fl/fl^*samples were generated in this way, from a total of 35 mice. The microglia were fixed with 1% formaldehyde at 20℃ for 8 min then quenched with 0.125 M Glycine. The nuclei were isolated and sonicated using a Diagenode disruptor for 20 cycles (30 seconds “ON”, 30 seconds “OFF”, high power). The chromatin fragments were enriched using 2 ul of anti-H3K9ac antibody (Millipore, 07-352) per reaction. The ChIPed DNA was purified and processed for library preparation. The ChIP DNA libraries were size selected using SPRIselect beads (Beckman Coulter, Cat.#B23317) and quantified using Agilent 2100 Bioanalyzer and Qubit (Invitrogen, Cat.#Q33238). The pooled libraries were then sent to Azenta Life Sciences for paired-end sequencing (2×150 bp) on Illumina HiSeq platform.

### Epigenetic data analysis

Raw fastq files were first checked for quality using Multiqc sequence analysis. Cutadapt (v.4.0) was used to cut adaptors (-*a* AGATCGGAAGAGCACACGTCTGAACTCCAGTC *-A* AGATCGGAAGAGCGTCGTGTAGGGAAAGAGTGT), and reads were aligned to the mouse genome (mm10) using bowtie2 (2.3.4.3)^60^. Subsequently, SAM files were sorted and filtered using Sambamba (v.0.8.2)^61^. We discarded unmapped and duplicate fragments. Sorted BAM files were indexed using samtools (v.1.15.1)^62^. Normalization for all BigWig files were carried out against effective genome size (2652783500 for mice H3K9ac ChIP-seq and ATAC-seq). Peak analysis was carried out with Macs2 (v.2.2.7)^63^ using stringent parameters (-f BAMPE --mfold 5 50 -p 0.001). A master-peak file was created, using the following settings for ATAC (-d 300 -c 7) and H3K9ac (d 300 -c 3). Raw counts were extracted from BAM files using master-peak file (**SI Table xxx**) and loaded into DESeq2^64^. Differentially accessible peaks were determined with adjustment for false discovery using benjamini hochberg method (Padj < 0.05). Differentially accessible peaks were annotated using HOMER (v4.4)^32^. BAM files were then converted to BigWig files and merged per experimental group for visualization using deeptools (v.3.5.0)^65^. Peak plots were constructed using deepotools with 1000bp extention from center of peak. Peaks were visualized using IGV (v.2.11.4), exported as .png, and edited in Adobe Illustrator. (For more details, see ‘Code availability’).

---

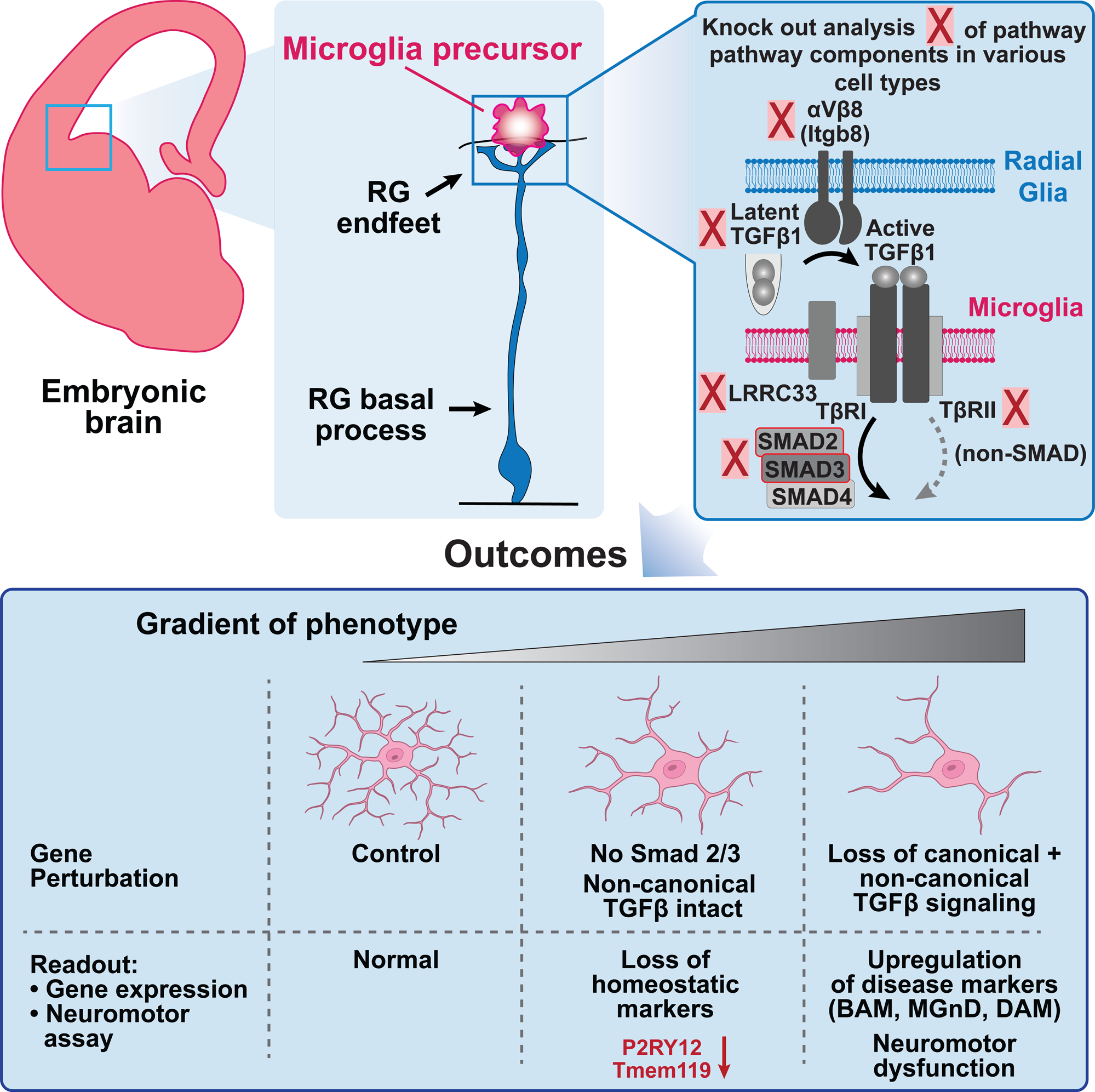

