## Supplemental Figures and Legends for "Radial glia promote microglial development through integrin α_V_β_8_-TGFβ1 signaling"

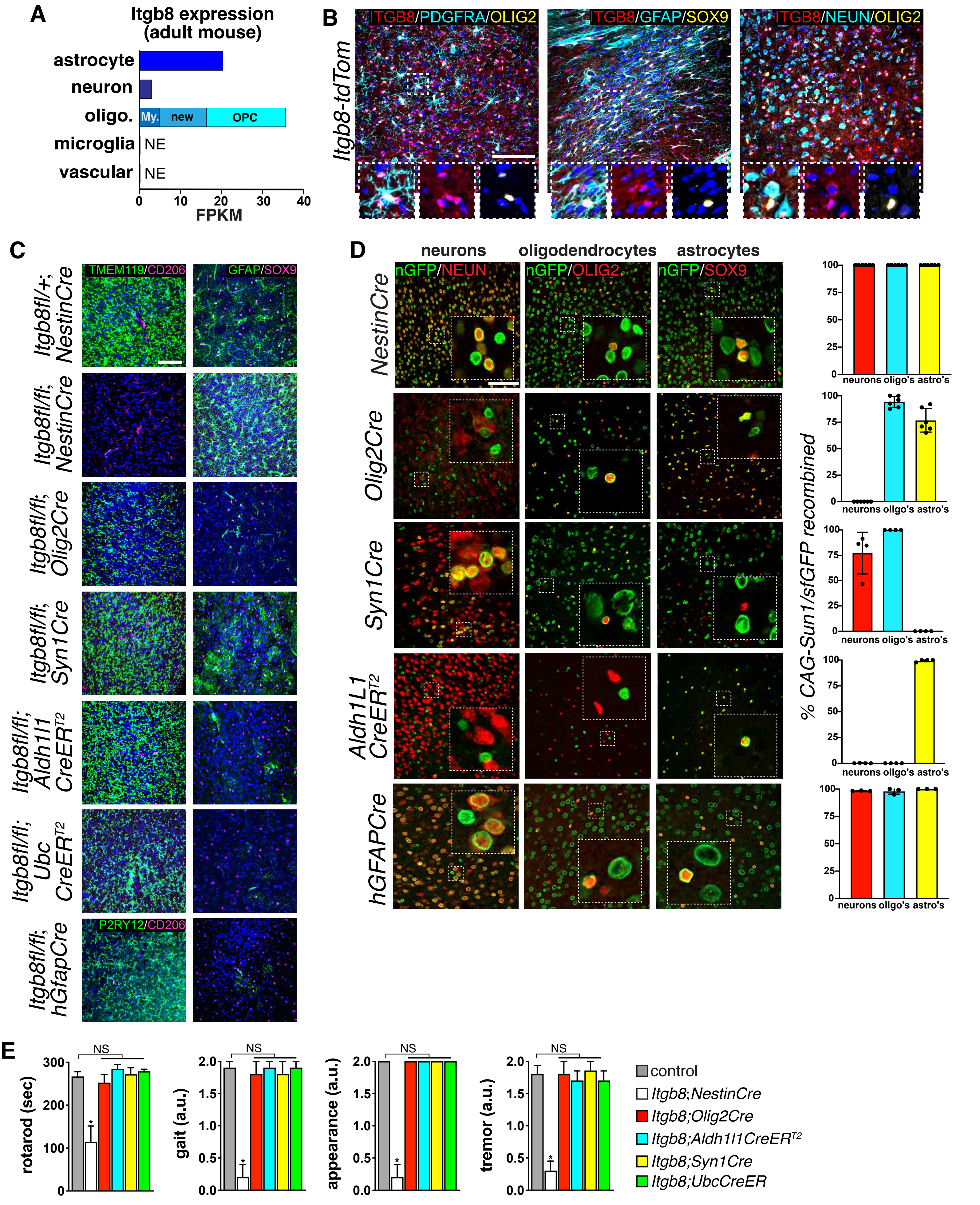


**Supplemental Figure 1. Deletion of *Itgb8* in adulthood does not disrupt microglial homeostasis.**

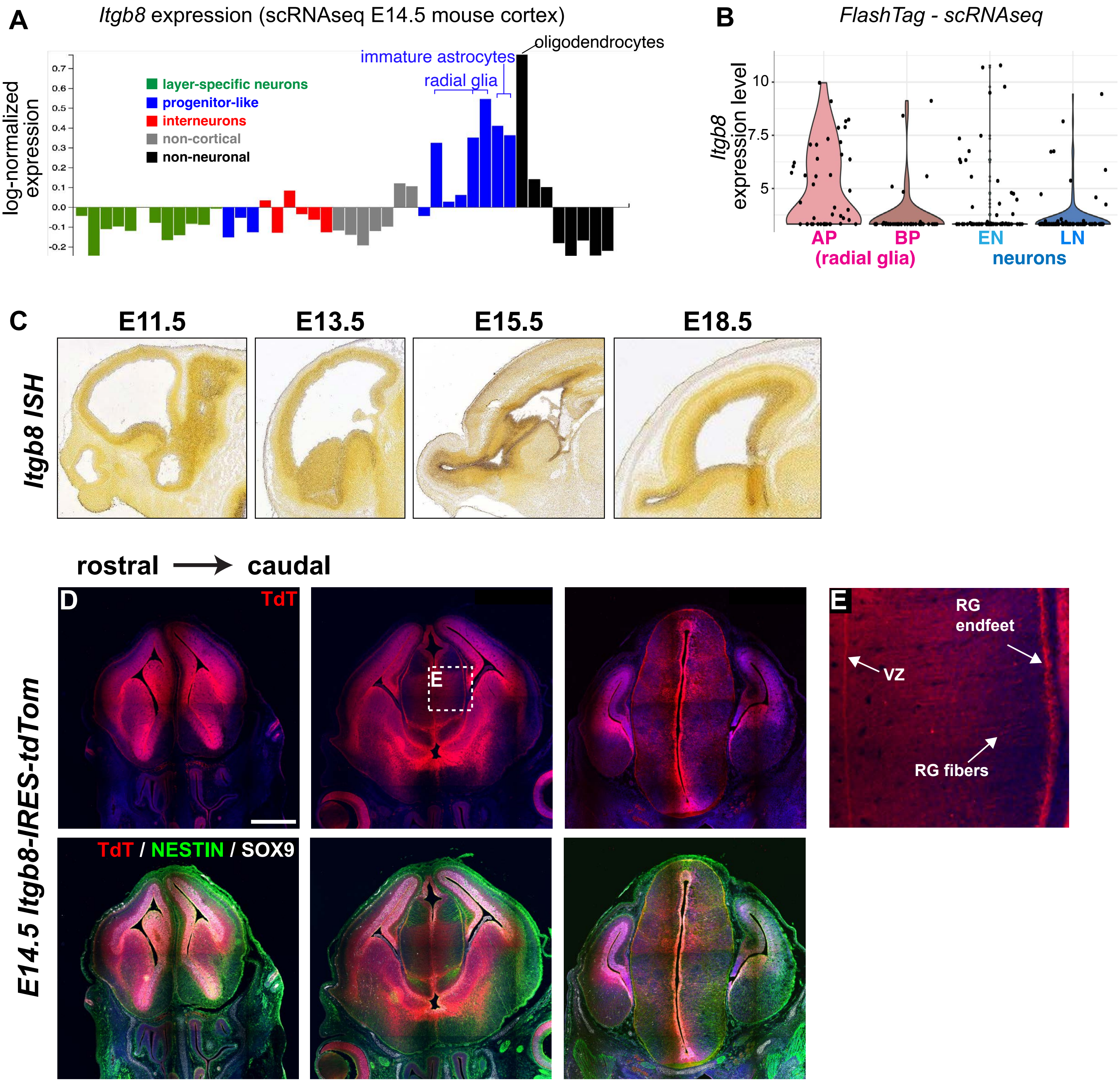


**Supplemental Figure 2: Analysis of cell-type specific *Itgb8* expression.**

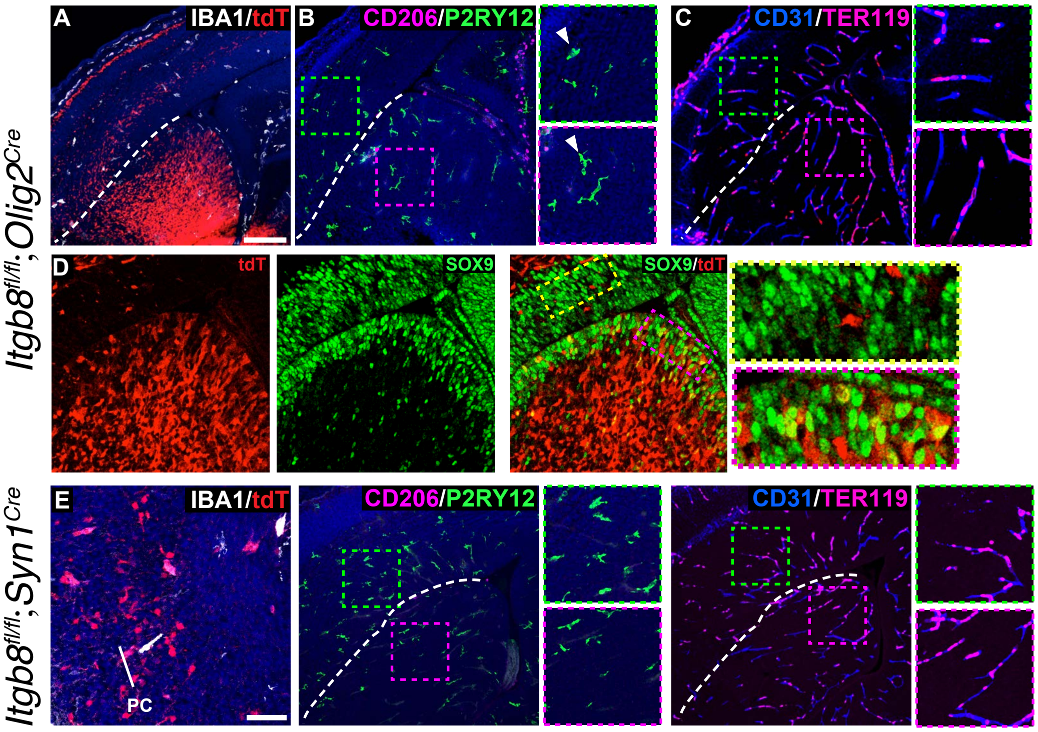


**Supplemental Figure 3: Cre-recombination and phenotypes in *Itgb8$^{fl/fl}$;Olig2$^{Cre}$* *and Itgb8fl/fl;Syn1$^{Cre}$* mice.**

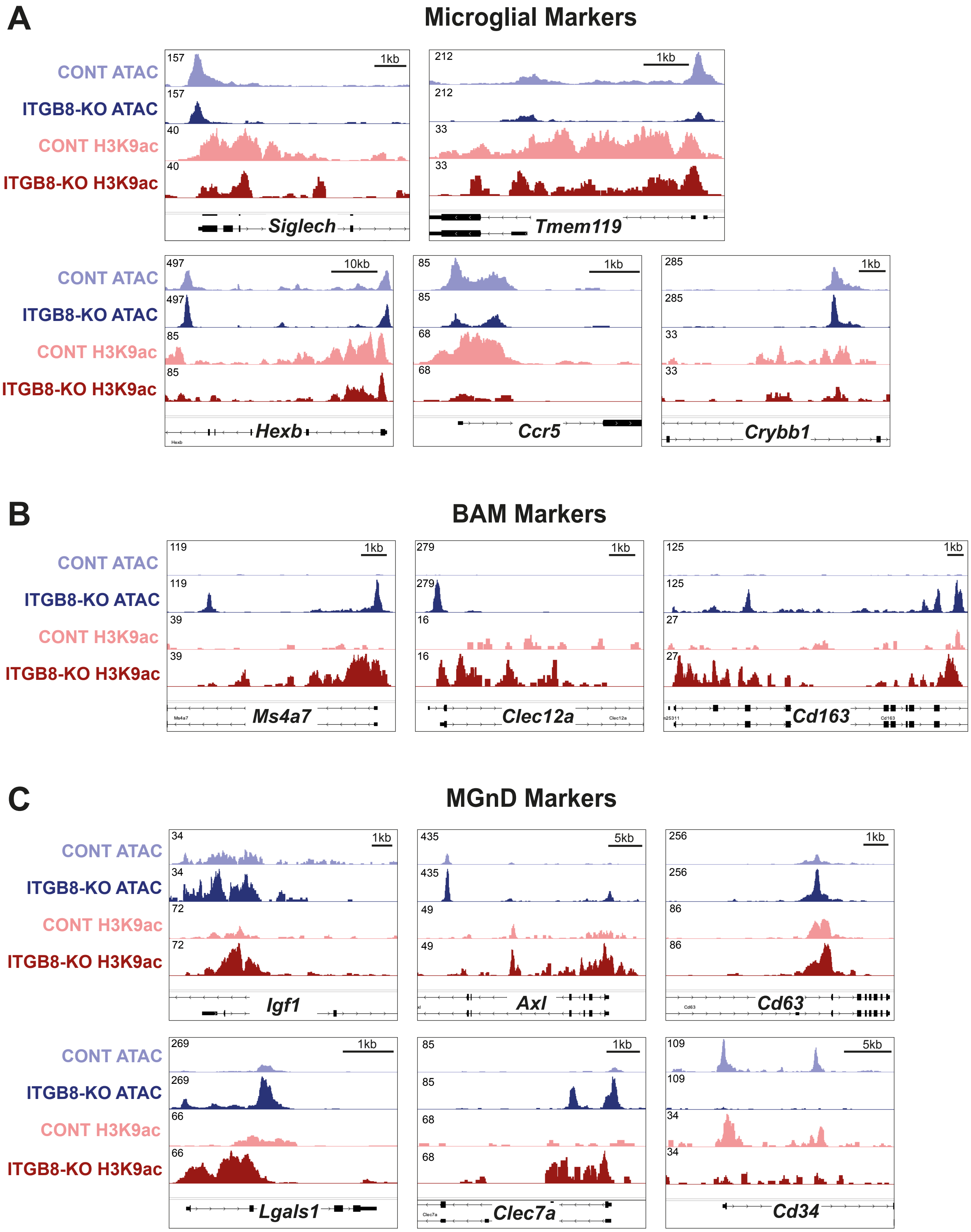


**Supplemental Figure 4:** ***Itgb8*** **deletion results in epigenetic changes in BAM and disease-associated macrophage gene bodies.**

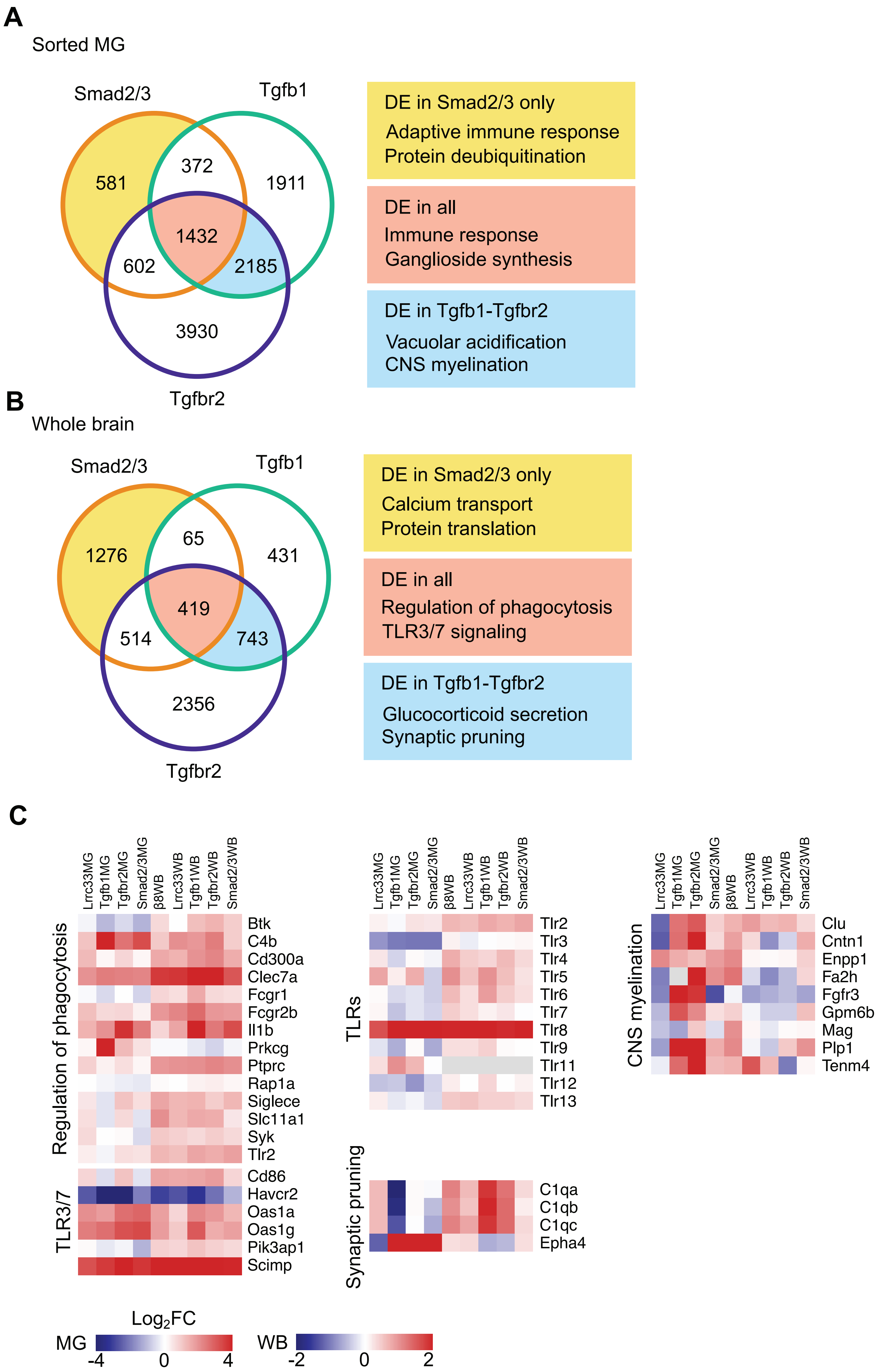


**Supplemental Figure 5: Gene ontology and transcriptional analysis of TGFβ mutant models.**

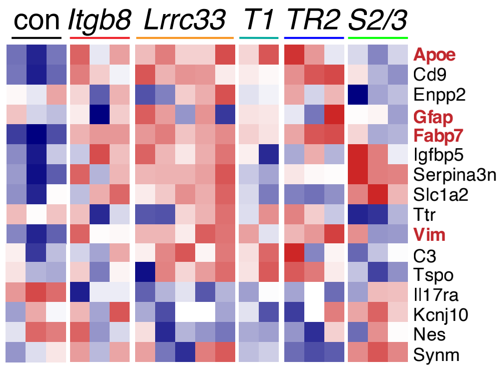


**Supplemental Figure 6: Astrocytosis-related gene expression in TGFβ pathway mutants.**

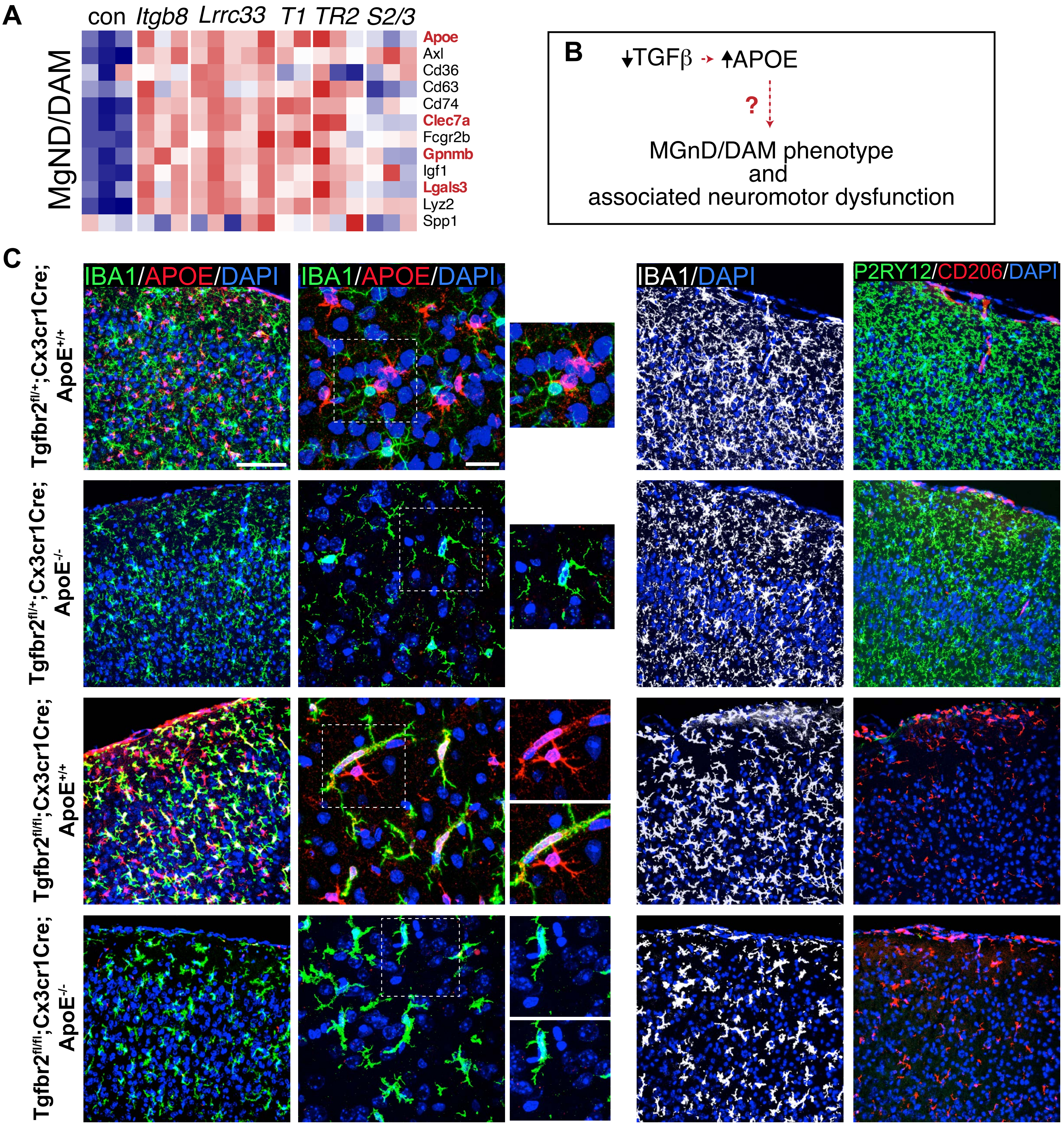


**Supplemental Figure 7: Epistatic analysis of *ApoE* contribution to the *TgfbR2* conditional mutant microglial phenotype.**

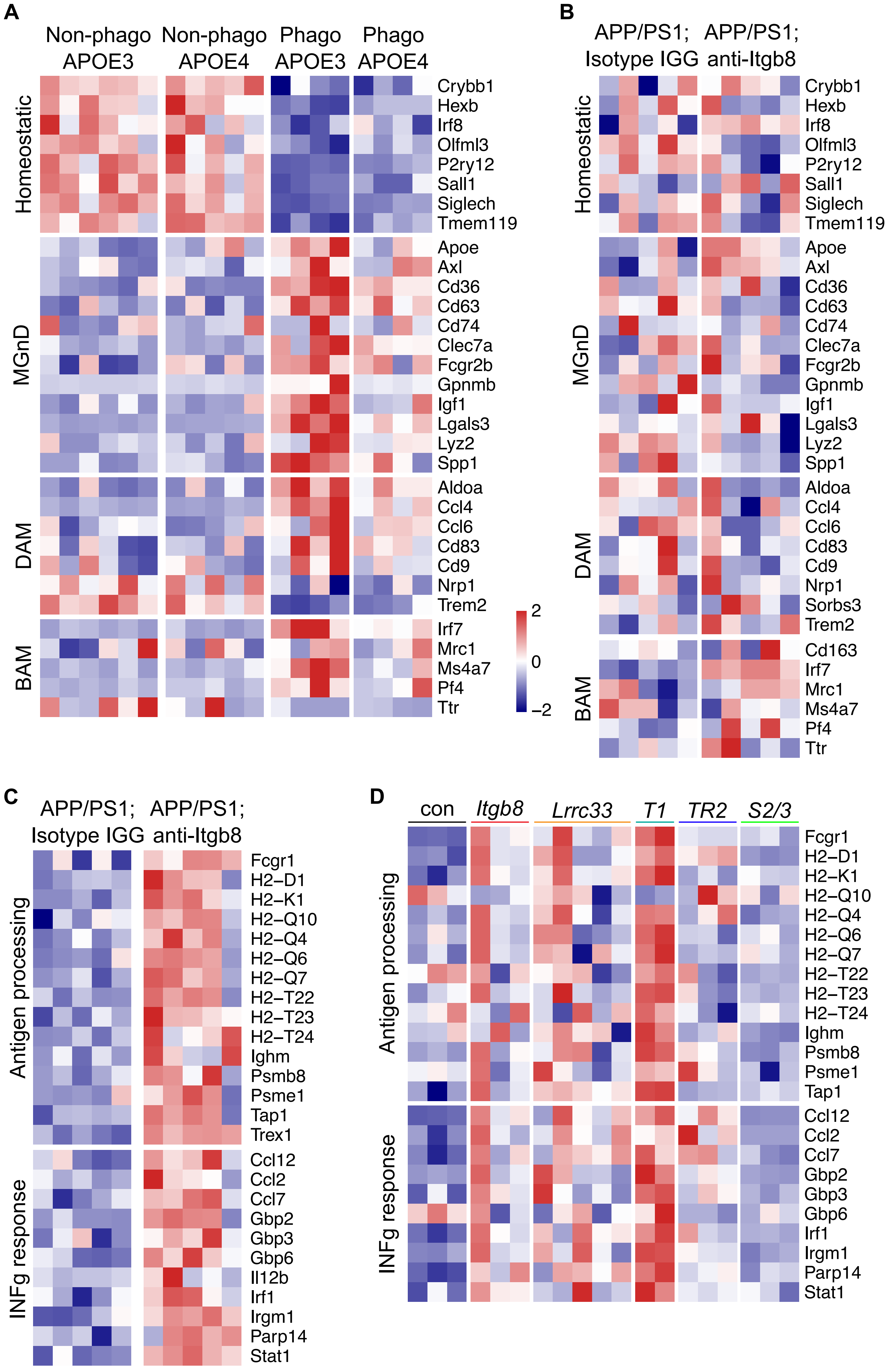


**Supplemental Figure 8: Gene expression analysis in APOE variant mouse models, ITGB8 blockade, and TGFβ mutant models.**

**Supplemental Figure Legends**

**Supplemental Figure 1. Deletion of *Itgb8* in adulthood does not disrupt microglial homeostasis.**

**A)** Expression analysis of *Itgb8* expression in adult astrocytes, neurons, oligodendrocytes, microglia, and endothelial cells[35]. **B)** *Itgb8* is expressed in oligodendrocytes (OLIG2+; PDGFRa+), OPCs (OLIG2+), astrocytes (SOX9, GFAP+/-), and neurons (NEUN+) in adulthood, as shown by immunohistochemical overlap with a *Itgb8-TdT* reporter line. **C)** Deletion of *Itgb8* using various cell-lineage restricted *Cre* and *CreER* mouse lines does not result in changes in microglial homeostasis or associated astrocyte activation, unlike deletion with $Nestin^{Cre}$. **D)** Cre-dependent nuclear GFP *Cre*-reporter line *SunTag* crossed to indicated *Cre* or *CreER* line, and brain sections from adult mice were stained for nuclear-localized markers of neurons (NEUN), oligodendrocytes and oligodendrocyte precursors (OPCs) (OLIG2), or astrocytes (SOX9); images taken from motor cortex. Cell marker colocalization with *CAG-Sun1/sfGFP* recombination to the right. 6 animals were quantified for $Nestin^{Cre}$ and $Olig2^{Cre}$, 4 for $Syn1^{Cre}$ and $Aldh1L1^{CreER}$ and 3 for $hGFAP^{Cre}$. 3 images per mouse were quantified from the cerebral cortex. **E)** Deletion of *Itgb8* using various cell-lineage restricted *Cre* and *CreER* mouse lines does not result in changes in motor behavior, unlike deletion with *Nestin-Cre*, which results in profound motor dysfunction. Scale bars in B,C and D=100$\mu$m.

**Supplemental Figure 2. Analysis of cell-type specific *Itgb8* expression.**

**A)** Single-cell RNA-seq analysis of *Itgb8* expression in the developing brain, from[17]. Individual columns represent clustered categories of cell types. **B)** *Itgb8* expression in apical and basal progenitors, early born and late born neurons, derived from Flash-Tag ScRNA-seq analysis in [18]. **C)** *Itgb8* in situ hybridization (ISH) at indicated time points taken from Allen Brain Atlas (https://mouse.brain-map.org). **D)** Analysis of $Itgb8^{tdT}$ expression in E14.5 mouse brain stained for radial glia nuclei (SOX9, white) and radial glia processes (NESTIN, green). **E)** Magnified image of the embryonic thalamus from **(D)**, showing $Itgb8^{tdT}$ expression in the ventricular progenitor zone, radial glia fibers and radial glia endfeet in the overlying meninges. Scale bar in D=500$\mu$m.

**Supplemental Figure 3. Cre-recombination and phenotypes in $Itgb8^{fl/fl};Olig2^{Cre}$ and $Itgb8^{fl/fl};Syn1^{Cre}$ mice.**

**A)** Coronal brain sections from E14.5 $Itgb8^{fl/fl};Olig2^{Cre}$ mice stained for tdT (Cre recombination, red) and microglia/macrophages (IBA1, white); **B)** microglia (P2RY12 in green, see arrowheads) and immature macrophages (CD206 in magenta); **C)** hemorrhage (red blood cells marked by TER119 in magenta and vascular endothelium marked by CD31 in blue); **D)** tdT (red) and apical progenitors (SOX9, green). Note lack of recombination of apical progenitors and also lack of microglia or vascular/hemorrhage phenotypes in these mice. **E)** Coronal brain sections from E14.5 $Itgb8^{fl/fl};Syn1^{Cre}$ stained for tdT (Cre recombination, red) and microglia/macrophages (IBA1, white); microglia (P2RY12 in green, see arrowheads) and immature macrophages (CD206 in magenta); hemorrhage (red blood cells marked by TER119 in magenta and vascular endothelium marked by CD31 in blue). PC=piriform cortex. Scale bar in A=150$\mu$m, scale bar in E= 50$\mu$m.

**Supplemental Figure 4. *Itgb8* deletion results in epigenetic changes in BAM and disease-associated macrophage gene bodies.**

**A-C)** Genome browser profiles for WT and Itgb8-KO ATAC-seq chromatin accessibility and histone H3K9ac ChIP-seq enrichment. Noted are prominent markers of **A)** microglia, **B)** borderzone macrophages and **C)** neurodegenerative disease associated (MGnD) microglia.

**Supplemental Figure 5. Gene ontology and transcriptional analysis of TGFβ mutant models.**

**A-B)** Gene ontology analysis of **A)** sorted microglia and **B)** bulk-Seq from different TGFβ mutant models. Putative genes involved in differential neuromotor phenotypes in TGFβ mutants were selected based on differential expression in distinct mouse models. Overrepresentation analysis was performed in each set separately as indicated by respective colors. **C)** Expression changes of genes driving enrichment of selected GO terms. Levels are shown as $Log_2$FoldChange from sorted microglia and whole brain from all analyzed mutants. We selected processes likely to be important in microglia-associated neuromotor impairments (i.e. regulation of phagocytosis, Toll-like receptor gene family and signaling, synaptic pruning, and myelination). Color scales are distinct for microglia and whole brain.

**Supplemental Figure 6. Astrocytosis-related gene expression in TGFβ pathway mutants.**

Comparative gene expression analysis of astrocytosis markers in various TGFβ pathway mutants reveals increased expression of astrocytosis markers in *Itgb8*, *Lrrc33*, *Tgfb1*, *Tgfbr2* conditional mutant models. In *Smad2/3* conditional mutants however, the astrocytosis markers (noted in red) *ApoE*, *Gfap*, *Fabp7* and *Vimectin* were not as highly upregulated.

**Supplemental Figure 7. Epistatic analysis of *ApoE* contribution to the *Tgfbr2* conditional mutant microglial phenotype.**

**A)** Heatmap of *ApoE* and other MGnD marker gene expression in TGFβ pathway mutants. **B)** Diagram describing potential pathway interaction between upregulated APOE in the *Tgfbr2* conditional mutant model and the downstream MGnD/DAM microglial phenotype. **B)** Analysis of effect of simultaneous deletion of *ApoE* and microglial *Tgfbr2*. No change was seen in $ApoE^{+/-}$; $TgfR2^{fl/+};Cx3cr1^{Cre}$ *versus* $ApoE^{-/-}$; $TgfR2^{fl/+};Cx3cr1^{Cre}$ in dysmature morphology, P2RY12 expression loss, CD206 upregulation. Scale bar in C=100$\mu$m. Scale bar in C=25$\mu$m for higher magnification image.

**Supplemental Figure 8. Gene expression analysis in APOE variant mouse models, ITGB8 blockade, and TGFβ mutant models.**

**A)** Expression of markers of homeostatic and neurodegeneration-associated microglia (MGnD), disease-associated microglia, and border-associated macrophages, in phagocytic and non-phagocytic Fcrls+ microglia from mice expressing APOE isoforms ε3 and ε4. **B)** Expression levels of the same markers in cortical microglia from APP/PS1 mice injected intracortically with an ITGB8-blocking antibody. **C)** Antigen processing and Interferon-γ related genes are upregulated in microglia after ITGB8-blocking antibody treatment in APP/PS1 mice. **D)** Levels of genes involved in those two processes in microglia from mice carrying inactivating mutations in TGFβ signaling components. Expression patterns are go mostly in opposite directions in *Smad2/3* condiitonal mutants compared to the other mutant lines.
